# Myeloid reprogramming by poly(I:C) recruits progenitor-exhausted CD8+ T cells and sensitizes rhabdoid tumors to PD-1 blockade

**DOI:** 10.64898/2025.12.29.696858

**Authors:** Valeria Manriquez, Sofia Cavada-Silva, Kévin Beccaria, Leticia L. Niborski, Amaury Leruste, Wilfrid Richer, Mathias Vandenbogaert, Zhi-yan Han, Christine Sedlik, Jordan Denizeau, Jérémy Mesple, Zoé Fusilier, L. Laëtitia Lesage, Jeremie Goldstein, Yohan Gerber-Ferder, Stéphanie Fitte-Duval, Federico Marziali, Rafael Mena-Osuna, Jimena Tosello-Boari, Rachida Bouarich-Bourimi, Maria Florencia Pacini, Yoann Missolo-Koussou, Mylene Bohec, Sylvain Baulande, Philippe Benaroch, Julie Helft, Hélène D. Moreau, Joshua J. Waterfall, Franck Bourdeaut, Eliane Piaggio

**Affiliations:** INSERM U932 Immunity and Cancer, Institut Curie Research Center, Paris, France; Department of Pediatric Neurosurgery-AP-HP, Necker Sick Kids Hospital, Paris, France; INSERM U1330 Children’s oncology research unit (CONCERT), Institut Curie Research Center, Paris, France; Department of Translational Research, Institut Curie Research Center, Paris, France; SIREDO: Care, innovation and research for children, adolescents and young adults with cancer, Institut Curie, Paris, France; INSERM U1016-CNRS UMR8104, Institut Cochin, Phagocytes et immunologie des cancers, Paris, France; PSL University, Institut Curie, ICGex Next-Generation Sequencing Platform, Paris, France; PSL University, Institut Curie, Single Cell initiative, Paris, France; Department of Pathology (PATHEX). Institut Curie, Paris, France; Paris-Cité University, Paris, France; PSL University, Paris, France

## Abstract

Immune exclusion remains a major barrier to effective immunotherapy in solid tumors. Given the abundance and plasticity of tumor-associated macrophages (TAMs) in many tumors, including pediatric tumors, we investigated whether TLR3 activation could reprogram them to facilitate immune access. Single-cell and spatial profiling in a mouse model of rhabdoid tumors showed that they are dominated by TLR3-expressing TAMs, whose depletion delays tumor growth. Treatment with the TLR3 agonist poly(I:C) promotes immune cell infiltration, including progenitor-exhausted CD8+ T cells, by multiple mechanisms including the reduction and reprogramming of immunosuppressive TAMs, promoting nitric-oxide production by peritumoral macrophages, and inducing CXCL9/10 production. Combined poly(I:C) and PD-1 blockade elicited durable, complete tumor rejection. Human macrophages from tumor biopsies showed conserved TLR3 responsiveness, underscoring translational potential. These findings uncover a mechanism by which TLR3-driven myeloid reprogramming transforms immune-excluded tumors into checkpoint-responsive ones, revealing a therapeutic path to overcome resistance to PD-1 blockade.

## Introduction

The efficacy of immune checkpoint blockade relies not only on the functional rescue of dysfunctional T cells but also on their ability to infiltrate tumors. In many solid cancers, limited immune infiltration remains a major barrier to effective immunotherapy^1,2^. Myeloid cells constitute the most abundant and functionally diverse immune population within tumors^3^, but whether they control immune access to the tumor microenvironment, and by which mechanisms, remains to be determined. Defining the mechanisms that control this infiltration is essential to expand the therapeutic reach of immunotherapy beyond currently responsive tumor types^4,5^.

SWI/SNF-deficient cancers, which account for nearly 20% of human malignancies^6,7^, display unique immunological features. Among them, rhabdoid tumors (RTs) are rare, highly aggressive pediatric cancers driven by the biallelic loss of SMARCB1, a core subunit of the SWI/SNF chromatin-remodeling complex^8–10^. Despite being genomically quiet and harboring one of the lowest tumor mutational burdens across cancers^11,12^, RTs exhibit signs of immune activation. Loss of SMARCB1 induces the re-expression of endogenous retroviruses and type I interferon signaling, resulting in a tumor microenvironment enriched in myeloid cells and clonally expanded CD8⁺ T cells expressing exhaustion markers such as PD-1, TIM-3, and LAG-3^13^. These observations challenge the notion that mutational load is the sole determinant of immunogenicity and position RTs as a paradigm to explore mechanisms of immune infiltration in genomically quiet but immunologically responsive tumors^14^. Consistently, both RT patients and RT-bearing mice show partial responses to PD1 blockade^13,15,16^, suggesting that additional mechanisms beyond T cell reinvigoration may limit durable tumor control.

Toll-like receptor 3 (TLR3) is a pattern-recognition receptor that detects double-stranded RNA (dsRNA) and triggers type I interferon signaling, bridging innate and adaptive immunity^17,18^. Polyinosinic:polycytidylic acid (poly(I:C)), a synthetic TLR3 agonist, has shown potent Immunostimulatory effects in preclinical models and is currently under clinical evaluation with minimal toxicity ^19–23^. Previous studies have proposed that poly(I:C) (PIC) enhances antitumor immunity by activating dendritic cells^24^ or modulating macrophage polarization^25–27^. However, the precise mechanisms through which TLR3 activation coordinates the interplay between innate and adaptive immunity within tumors remain unclear. Considering the abundance and heterogeneity of myeloid cells in RTs, and their association with immune suppression, we hypothesized that reprogramming the myeloid compartment could enhance antitumor immunity when combined with T cell reinvigoration.

Using scRNA-seq, TCR-seq, flow cytometry, and multiplexed imaging, we dissected how poly(I:C) treatment reprograms the myeloid compartment in RTs, promoting the recruitment and activation of progenitor-exhausted CD8⁺ T cells. This immune remodeling sensitizes tumors to PD-1 blockade, and the combination of poly(I:C) with anti-PD-1 results in a robust antitumor response, highlighting the relevance of TLR3-driven myeloid reprogramming as a strategy to enhance immune checkpoint therapy.

## Results

### TLR3 activation by poly(I:C) targets myeloid cells and controls rhabdoid tumor growth

Tumor-associated macrophages and other myeloid populations have been consistently linked to immune suppression and tumor progression across solid cancers^28,29^. To gain mechanistic insight into their contribution to rhabdoid tumors (RTs) and identify actionable targets for therapeutic reprogramming, we profiled the immune compartment of two large syngeneic R26S2 RTs (∼1500 mm³) by single-cell RNA sequencing (scRNA-seq) (**Figure 1A**). This model, derived from *Smarcb1^flox/flox; Rosa26-CreERT2* mice^30^, generates tumors in the flank of immunocompetent C57BL/6J hosts that faithfully recapitulate extracranial (ECRT) and MYC human RTs^13^. To define the major immune populations present in these tumors, we first analyzed FACS-sorted live cells using scRNAseq from 10X Genomics. After filtering the data to exclude dead, low-quality cells and stroma cells (without *Ptprc* expression), the UMAP visualization of the remaining 3,290 immune cells revealed eleven clusters, which were re-classified in seven major clusters (Figure S1A-B): monocytes and macrophages (MoMa, characterized by the expression of *Csf1r, Adgre1, Mrc1*), T cells (expressing *Cd3g, Cd3e, Cd28*), granulocytes (expressing *S100a9, Mmp9, Csf3r*), cycling cells (expressing *Stmn1, Ube2c, Top2a*), natural killer cells (NK, expressing *Ncr1, Gzma, Klrb1c*), dendritic cells (DCs, expressing *Batf3, Flt3, Xcr1*), and B cells (expressing *Cd79b, Ms4a1, Cd19*) (**Figure 1B-C**, S1D and Table S2). Quantitative analysis of the relative proportion of these populations per mice revealed that approximately 61% of the cells (n=2002) were classified as MoMa, identifying them as the most significant infiltrating subpopulation in RTs (**Figure 1D** and S1C). To determine the effect of MoMa on RT growth, we depleted this population by administering human diphtheria toxin (hDT) intraperitoneally to CD64^hDTR^ mice with palpable tumors (30-100 mm³ in volume). MoMa depletion led to a significant reduction in tumor growth, highlighting the critical pro-tumoral role of these cells and their potential as a clinical target (**Figure 1E** and S1E).

**Figure 1.**
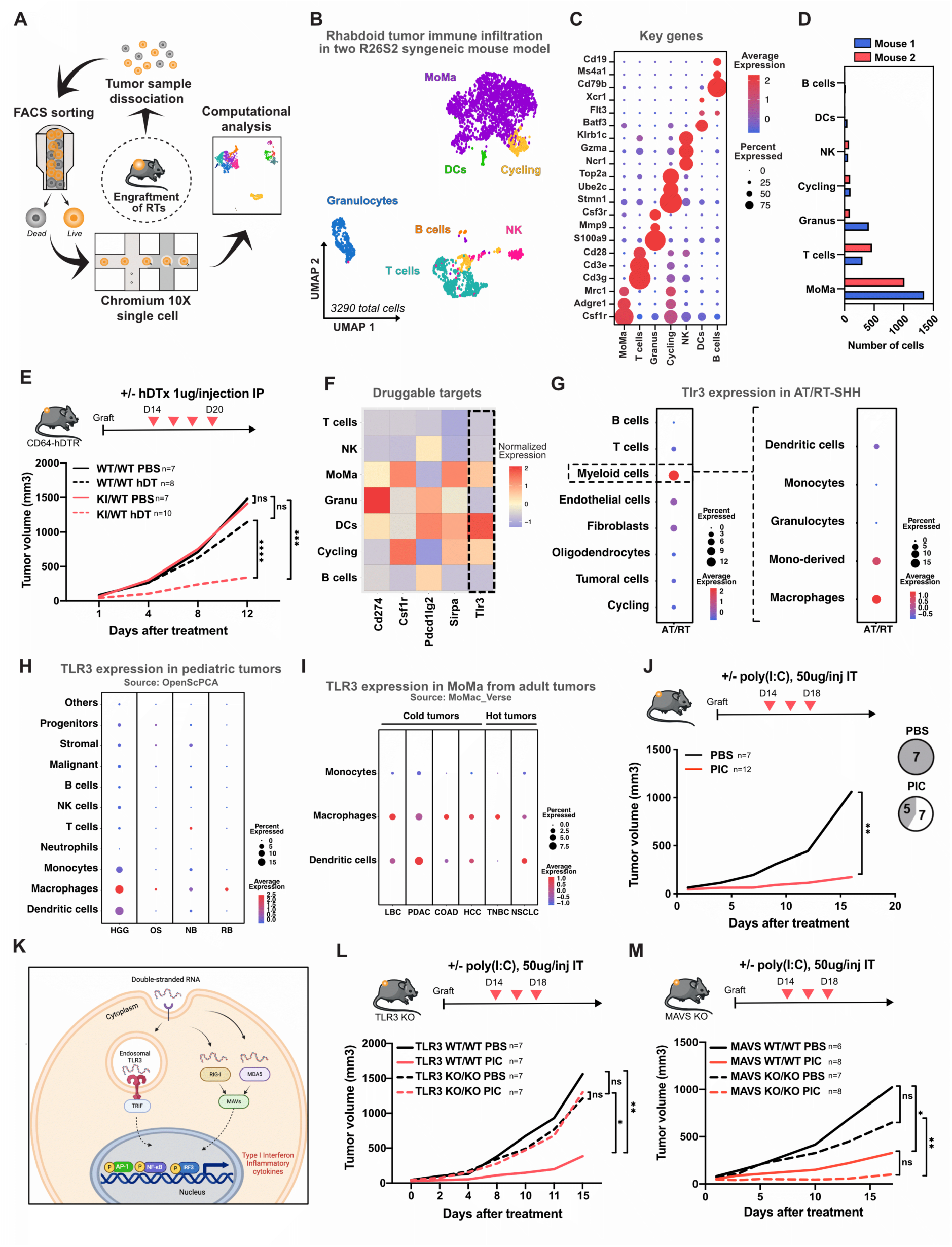
PIC targeting of TAMs and DCs via TL3 activation boosts the anti-tumor immune response. **A)** Schematic representation of the experimental workflow. **B)** UMAP visualization of 3290 CD45+ cells from two mouse RTs (volume ∼ 1500mm³) showing seven distinct clusters; Monocyte and macrophages (MoMa), T cells, Granulocytes (Granus), cycling cells, Natural Killer cells (NK), Dendritic cells (DCs) and B cells. **C)** Dot plot illustrating the scaled expression of three key genes per cluster, with dot size indicating the percentage of cells in each cluster expressing the corresponding gene. **D)** Stacked bar displaying the average relative proportion of each cluster per sample. **E)** RT-bearing CD64hDTR WT and KO mice were treated intraperitoneally with four doses of human diphtheria toxin (hDT, 1 μg per injection) or PBS (control) on alternate days, beginning when tumor volumes ranged from 30 to 100 mm³. Tumor growth curves are presented as the mean of at least 7 mice per group. Data shown are representative of one of two independent experiments. **F)** Heatmap representation of the relative mean expression of selected myeloid druggable targets (*Cd274* (PD-L1), *Csf1r*, *Pdcd1lg2* (PD-L2), *Sirpa*, and *Tlr3*) across different clusters. **G)** Dotplot illustrating TLR3 expression across cell populations (left) and within myeloid subsets (right) in human ATRT-SHH tumor. **H)** Dotplot illustrating TLR3 expression in pediatric tumors from *Open single cell pediatric cancer atlas* (OpenSCPCA). HGG: high-grade glioma, OS: osteosarcoma, NB: neuroblastoma and RB: retinoblastoma. **I)** Dotplot illustrating TLR3 expression in monocytes/macrophages (MoMa) from adult tumor scRNAseq datasets from *MoMa_Verse*. LBC: Luminal breast cancer, PDAC: Pancreatic ductal adenocarcinoma, COAD: Colon Adenocarcinoma, HCC: Hepatocellular Carcinoma, TNBC: Triple-negative breast cancer and NSCLC: Non-Small Cell Lung Cancer. **J-L-M)** RT-bearing mice were treated intratumorally with three doses of PIC (50ug/injection) or PBS (control) every other day, starting when tumor volume was between 30-10mm^3^. **J)** Tumor growth curves of C57BL/6J RT-bearing mice treated with PIC or PBS (control). The number of mice achieving complete tumor control out of the total treated is indicated in white on the circle. Data are presented as the mean of at least 7 mice per group, and are representative of one of two independent experiments. **K)** Diagram illustrating the recognition pathways of double-stranded RNA. Figure created using Biorender (https://biorender.com/ ). **L)** Tumor growth curves of TLR3 WT or KO tumor-bearing mice treated with PIC or PBS. Data are presented as the mean of at least 7 mice per group, and are representative of one of two independent experiments. **M)** Tumor growth curves of MAVs WT or KO tumor-bearing mice treated with PIC or PBS. Data are presented as the mean of at least 6 mice per group, and are representative of one of two independent experiments. **E-J-L-M)** Significance is calculated using Mann-Whitney test. *p<0,05, **p<0,01.

Several therapeutic strategies aimed at modulating the number and function of myeloid cells are currently under preclinical and clinical evaluation. Using scRNAseq data from the immune infiltrate, we analyzed the expression of several candidates, including *Cd274* (PD-L1), *Csf1r*, *Pdcd1lg2* (PD-L2), *Sirpa*, and *Tlr3*, across the different immune cell clusters. Our analysis revealed that most of these molecules, except PD-L2, were predominantly expressed by MoMa (**Figure 1F**). We focused on TLR3, selectively expressed by DCs and macrophages, because previous studies have shown that TLR3 ligands can induce IFN-beta-dependent tumor regression^31^ and conventional dendritic cell activation, essential for priming T cells in draining lymph nodes (LNs)^24,32^. To explore the translational potential of TLR3 targeting in human tumors, and more specifically in pediatric tumors, we analyzed its expression across multiple datasets. scRNA-seq of human RTs confirmed that TLR3 was preferentially expressed in myeloid populations, including macrophages and DCs (**Figure 1G**, S1F and Table S2). Similar TLR3 expression was observed in other pediatric tumors, high-grade glioma, osteosarcoma, neuroblastoma, and retinoblastoma (**Figure 1H**), as well as in adult malignancies, including luminal and triple-negative breast cancer, pancreatic, colorectal, hepatic, and non-small cell lung cancer (**Figure 1I**). Together, these data demonstrate that TLR3 expression within the myeloid compartment is conserved across diverse tumor types, underscoring the translational relevance of targeting this pathway and supporting RT mouse tumors as a powerful model to explore its therapeutic potential.

To test the therapeutic efficacy of targeting TLR3 in RT, we administered poly(I:C) (PIC), a synthetic TLR3 ligand composed of double-stranded RNA, intratumorally to RT-bearing mice. This treatment induced a significant reduction in tumor growth, with complete tumor regression observed in 7 out of 12 treated mice (**Figure 1J** and S1G). Given that PIC is currently under investigation for the treatment of some pediatric and non-pediatric solid tumors, where it has been administered intramuscularly with a favorable safety profile^21,23,33,34^, we sought to determine whether the route of administration influenced its antitumor effect. We observed that doubling the dose of PIC delivered intramuscularly resulted in the same level of efficacy as intratumoral injection, without inducing signs of toxicity (Figure S1I). Similar findings were also reported in mice with melanoma^24^, further supporting the broad potential of this approach.

It has been reported that double-stranded RNA (dsRNA) can be recognized by two types of pathogen recognition receptors (PRRs): TLR3 and RIG-I-like receptors (RLRs), with the latter activating a MAVs-associated signaling pathway^35^ (**Figure 1K**). To discern whether the anti-tumor effect mediated by PIC occurs via TLR3 or RLRs, TLR3 and MAVS knockout (KO) mice were treated with PIC. The anti-tumor efficacy of PIC was significantly impaired only in TLR3 KO, but not in MAVs KO mice, suggesting that PIC predominantly drives the anti-tumor response mainly through TLR3, rather than through RLRs (**Figure 1L-M**, and S1J-K). Importantly, in this model, the tumor fragments engrafted into TLR3 KO mice were derived from the wild type R26S2 syngeneic mouse model, meaning that the tumor cells themselves expressed TLR3 (Figure S1H), and suggesting that the anti-tumor effect of PIC is mainly due to its modulation of the host immune cells rather than a direct effect on the tumor cells. Overall, our results demonstrate that targeting immunosuppressive myeloid cells via PIC-mediated TLR3 activation significantly reduces tumor growth by modulating the host immune cells.

### PIC enhances granulocyte and monocyte tumor-infiltration while decreasing DCs and macrophages

To understand the effects of PIC on myeloid cells, RT-bearing mice were treated with a single intratumoral injection of PIC, and tumors were collected 16 hours later for analysis by scRNA-seq, flow cytometry (FACS), and immunofluorescence. This time point was chosen to assess myeloid cell activation while minimizing potential confounding effects from secondary T cell activation **(Figure 2A)**. We performed scRNA sequencing of FACS-sorted myeloid cells (CD45⁺ TCRβ⁻ NK1.1⁻ CD19⁻) from tumors from two PBS and two PIC-treated mice. UMAP visualization of the 21,936 total cells revealed the presence of three major cell clusters: dendritic cells (DCs, expressing *Flt3, Batf3*, and *Xcr1*), monocytes/macrophages (MoMA, expressing *Lyz2*, *Apoe* and *Csf1r*), and granulocytes (expressing *Csf3r, Mmp9* and *S100a9*) **(Figure 2B-C and S2A-C)**. Each cluster displayed unique genes or gene signatures previously described in the literature, further confirming their identity **(Figure 2D,S2E and Table S3)**. Density plots revealed an accumulation of granulocytes, a reduced presence of DCs, and a redistribution of MoMA following PIC treatment **(Figure 2E)**. These patterns were consistent across all four analyzed samples **(Figure S2D)**. FACS-based quantification further confirmed these trends, demonstrating that PIC treatment led to a significant enrichment of granulocytes and monocytes, while tumor-associated macrophages (TAMs) and DCs were significantly reduced, both in percentages and absolute numbers per mg of tumor **(Figure 2F and S2F)**.

**Figure 2:**
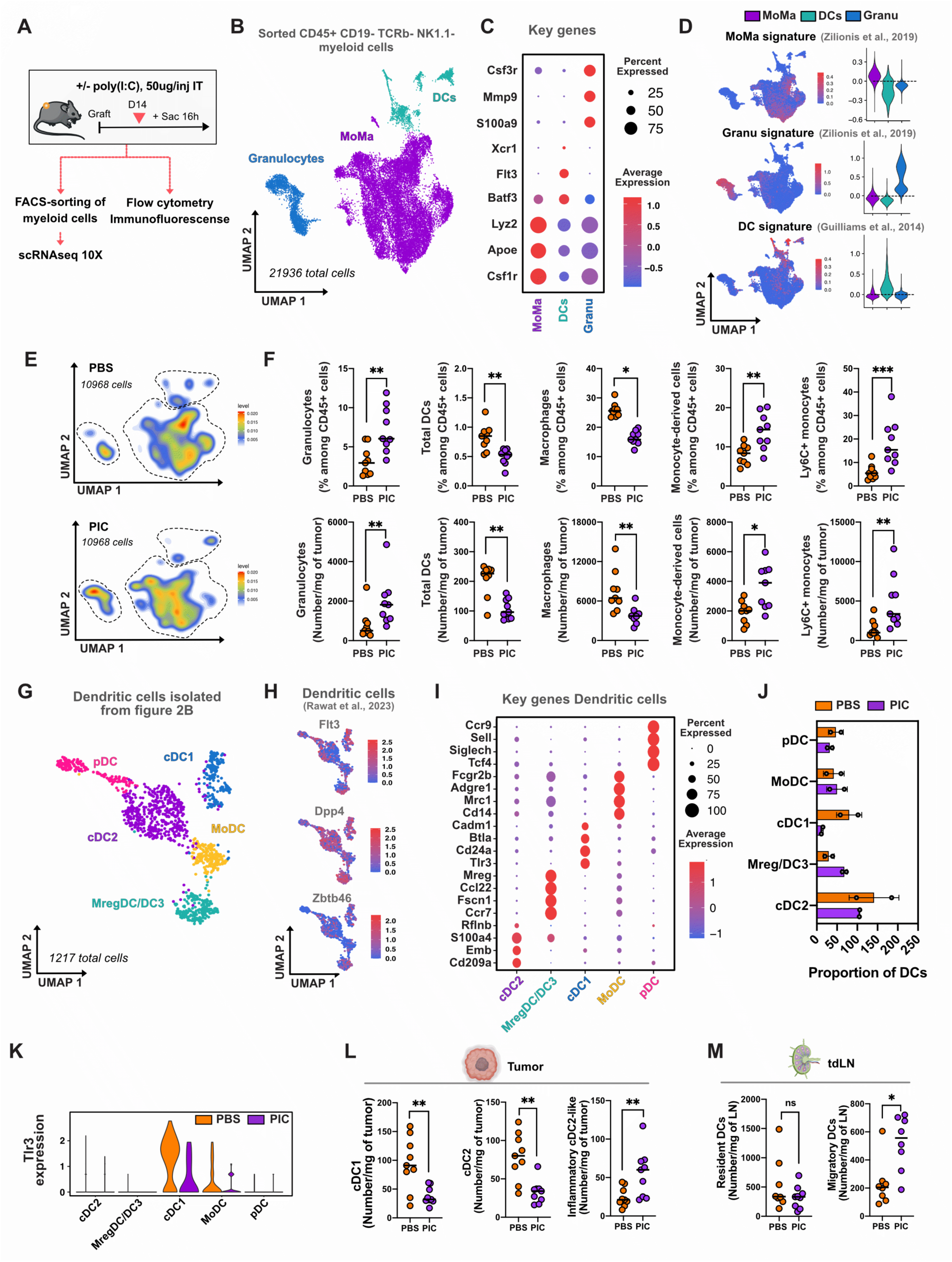
Differential effect of PIC on myeloid cell subsets. (**A**) Schematic representation of the experimental workflow. RT-bearing mice received a single intratumoral (IT) injection of PIC, and tumors were collected 16 hours later for analysis by scRNA-seq, flow cytometry (FACS), and immunofluorescence. **(B)** UMAP visualization of 21,936 CD45⁺TCRβ⁻NK1.1⁻CD19⁻ myeloid cells from two PBS-treated and two PIC-treated mice, revealing three major clusters: dendritic cells (DCs), monocytes/macrophages (MoMA), and granulocytes (Granus). **(C)** Dot plot showing the scaled expression of key genes per cluster. Dot size represents the percentage of cells within each cluster expressing the corresponding gene. **(D)** Enrichment of MoMA, granulocyte, and DC signatures. Left: UMAP visualization showing the expression of each signature within the same UMAP plot as in (B). Right: Violin plots displaying the distribution of signature expression across clusters. **(E)** Density plots depicting the spatial distribution of 10,968 myeloid cells per condition. Top: PBS-treated tumors. Bottom: PIC-treated tumors. **(F)** FACS-based quantification of granulocytes, total DCs, macrophages, monocyte-derived cells, and Ly6C⁺ monocytes. Top, frequency of each population among CD45+ cells in PBS- and PIC-treated tumors. Bottom, number of cells per mg of tumor in PBS- and PIC-treated tumors. Data shown are representative of two independent experiments, with n=9 mice per condition in the experiment presented and n=6 mice per condition in the replicate experiment, both yielding consisting results. **(G)** UMAP visualization of 1,217 dendritic cells isolated from (B), identifying five distinct clusters: cDC2, Mreg/DC3, cDC1, MoDC, and pDC. **(H)** UMAP visualization of dendritic cell-specific gene expression; Flt3, Dpp4, and Zbtb46. **(I)** Dot plot showing the scaled expression of key genes per DC cluster. Dot size represents the percentage of cells within each cluster expressing the corresponding gene. **(J)** Bar graph displaying the average proportion of each DC cluster. **(K)** Violin plot comparing Tlr3 expression levels across DC subsets in PBS- and PIC-treated tumors. **(L)** FACS -based quantification of conventional DCs in RTs. Data shown are representative of two independent experiments, with at least n=8 mice per condition in the experiment presented and n=6 mice per condition in the replicate experiment, both yielding consisting results. **(M)** FACS-based quantification of resident and migratory DCs in tumor-draining lymph nodes (tdLN). Data shown are representative of two independent experiments, with at least n=8 mice per condition in the experiment presented and n=6 mice per condition in the replicate experiment, both yielding consisting results. **(F-L-M)** Mann-Whitney test. *p<0.05, **p<0.01, ***p<0.001.

### PIC enhances DC migration to lymph nodes

To further investigate the impact of PIC on DCs, we bioinformatically isolated the DC cluster from Figure 2B, which contained 1,217 cells, and reintegrated it for detailed analysis. UMAP visualization revealed five distinct DC clusters **(Figure 2G and S2G)**, broadly characterized by the expression of *Flt3, Dpp4* and *Zbtb46* **(Figure 2H)** as previously described^36,37^. Based on unsupervised clustering and gene signatures from the literature, we identified five DC clusters **(Figure 2G-I, S2H, and Table S3-S7):** conventional type 2 dendritic cells (cDC2, expressing *Rflnb, S100a4, Emb,* and *Cd209)*, Mreg/DC3 (expressing *Mreg, Ccl22, Fscn1,* and *Ccr7),* conventional type 1 dendritic cells (cDC1, expressing *Cadm1, Btla, Cd24a,* and *Tlr3)*, monocyte-derived dendritic cells (MoDC, expressing *Fcgr2b, Adgre1, Mrc1,* and *Cd14),* and plasmacytoid dendritic cells (pDC, expressing *Ccr9, Sell, Siglech,* and *Tcf4)* **(Figure 2G,2I)**.

Quantitative analysis of the proportion of each cluster revealed a trend towards decreased cDC1 and increased Mreg/DC3 in tumors following PIC treatment **(Figure 2J)**. Interestingly, cDC1 exhibited the highest *Tlr3* expression in our dataset **(Figure 2K)**, aligning with previous findings that TLR3 ligands activate cDC1, promoting their migration to lymph nodes to prime T cells and enhance anti-PD1 responses^24,32^. To assess whether PIC treatment affected DC migration, we performed FACS analysis, which confirm a reduction of cDC1 and cDC2 in tumors and a corresponding increase in migratory DCs in tumor-draining lymph nodes **(Figure 2L-M, S2I)**. Together, these findings suggest that PIC promotes DC migration to lymph nodes, potentially enhancing antigen presentation and T cell activation.

### PIC reduces pro-tumoral macrophages and reprograms remaining TAMs to enhance anti-tumor immunity

To evaluate the impact of PIC on TAMs, we bioinformatically isolated and reintegrated the MoMa clusters from Figure 2B for detailed analysis. UMAP visualization identified macrophages (*Cx3cr1, Pdgfc, Cxcl9, IFN, Mreg*, and *cycling*), monocytes (*Ednrb, Rpsb, Sell*, and *non-classical*), and monocyte-derived cells (MD and MD-DC) **(Figure 3A, S3A-C)**. Macrophages were characterized by the expression of *Csf1r* and *Apoe* but lacked *Ly6c2* expression. Monocyte-derived cells expressed *Csf1r*, *Ly6c2*, *H2-Aa*, *Apoe*, and *Itgax*, whereas monocytes were identified by *Ly6c2* and *Ace* but exhibited low or no expression of *H2-Aa*, *Apoe*, or *Itgax* **(Figure 3B)**. This classification was further validated by differential gene expression (DGE) analysis, which confirmed the enrichment of characteristic genes in each population: *C1qa, Cxcl9, Folr2*, and *Trem2* in macrophages (Mac), *Ly6c2, S100a11, Ifitm6, Napsa*, and *S100a8* in monocytes (Mono), and *H2-Aa, H2-Ab1, Cd74, Itgax*, and *Mgl2* in MD cells **(Figure 3C)**. Furthermore, trajectory analysis supported the classification of these populations, revealing a differentiation continuum from monocytes to MD cells and ultimately to macrophages **(Figure 3D)**.

**Figure 3:**
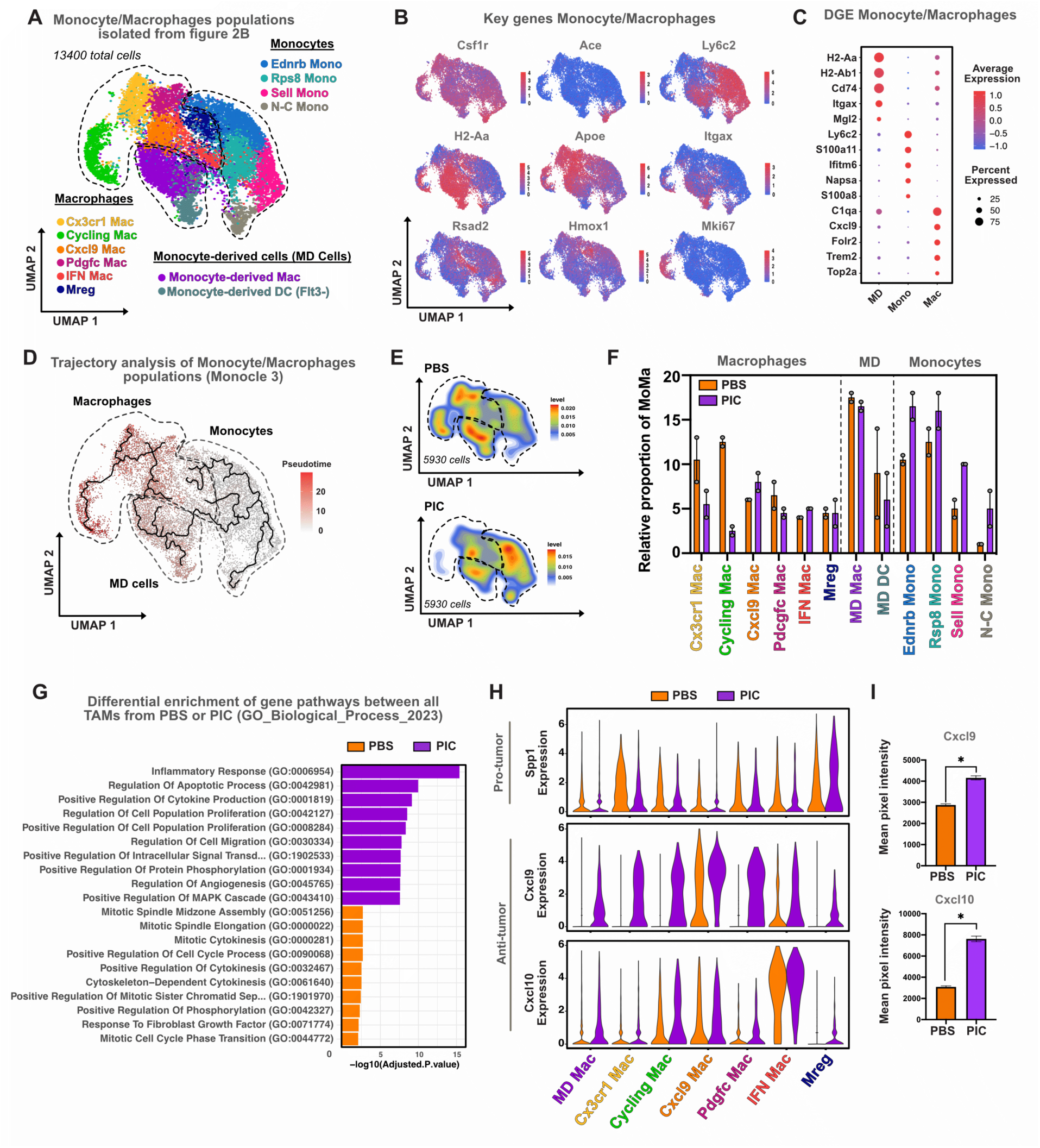
PIC treatment reprograms TAMs towards an anti-tumoral phenotype. (**A**) UMAP representation of the monocyte/macrophage (MoMa) compartment, isolated from Figure 2B (13,400 cells). Populations were classified into monocytes (Ednrb, Rpsb, Sell, and non-classical), monocyte-derived cells (MD cells), and macrophages (Cx3cr1, Pdgfc, Cxcl9, IFN, Mreg, and cycling macrophages). **(B)** Feature plots showing key marker genes distinguishing monocytes, monocyte-derived cells, and macrophages. **(C)** Dot plot of differentially expressed genes (DEGs) highlighting marker genes for macrophages (Mac), monocytes (Mono), and monocyte-derived cells (MD). **(D)** Trajectory analysis of monocyte/macrophage populations using Monocle 3. Pseudotime coloring indicates progression along a differentiation trajectory. **(E)** UMAP density plots comparing the distribution of MoMa populations between PBS and PIC-treated tumors. Top: PBS-treated tumors. Bottom: PIC-treated tumors. 5930 cells per condition. **(F)** Bar graph displaying the relative proportion of monocyte, monocyte-derived and macrophage subsets in PBS and PIC conditions. **(G)** Gene set enrichment analysis (GSEA) comparing gene pathways in macrophages from PBS and PIC treated tumors. Analysis was performed using GO biological process 2023 database. For each comparison, DEGs with an avg_log2FC > 0.25 and an adjusted p-value (p_val_adj) <0.05 were selected as input for the analysis. **(H)** Violin plots depicting the expression of *Spp1*, *Cxcl9* and *Cxcl10* across macrophage subsets in PBS and PIC-treated tumors. **(I)** Quantification of Cxcl9 and Cxcl10 protein levels in tumor supernatants from PBS and PIC-treated mice using the Proteome Profiler Mouse XL Cytokine Array (R&D System).

We first assessed the quantitative changes in cell populations following PIC treatment using a UMAP density plot, which revealed a global shift in the MoMa compartment, characterized by an increase in monocytes and a decrease in macrophages **(Figure 3E)**. Cluster-specific analysis confirmed that all monocyte subsets increased following PIC treatment, whereas monocyte-derived cells remained largely unchanged. Among macrophages, a tendency to reduction was observed in the most mature populations: the cycling, Cx3cr1, and Pdgfc subsets. Myeloid regulatory cells (Mreg) remained unaffected and the high TLR3-expressing Cxcl9 and IFN macrophages (**Figure S3E**) tended to increase **(Figure 3F)**. Given that TLR3 expression was primarily detected in macrophages, but not in monocytes (**Figure S3D**), we investigated PIC effect on macrophages phenotypes. Gene set enrichment analysis of total macrophages revealed pathways associated with cytokinesis, tissue remodeling, and homeostasis maintenance in untreated tumors, while PIC treatment enriched pathways related to cytokine production, cell migration, and immune signaling, indicating a shift towards a pro-inflammatory phenotype (**Figure 3G**). Furthermore, we analyzed the expression of *Cxcl9* and *Cxcl10*, previously linked to anti-tumoral functions and improved patient survival; and of *Spp1*, previously associated with pro-tumoral activity and poorer prognosis^38^. In the PBS condition, *Spp1* showed the highest expression in Cx3cr1 and Mreg macrophages, while *Cxcl9* and *Cxcl10* were most highly expressed in Cxcl9 and IFN macrophages, respectively. Upon PIC administration, *Spp1* expression was reduced across all clusters except Mreg, while *Cxcl9* and *Cxcl10* expression increased in all macrophage subpopulations **(Figure 3H)**. To assess if this upregulation correlated with protein levels, tumor supernatants were analyzed, confirming a significant increase in both chemokines **(Figure 3I)**.

Collectively, these results suggest that PIC treatment reduces Cx3cr1 pro-tumoral macrophages and reprograms the remaining macrophages towards an anti-tumoral phenotype, marked by increased Cxcl9 and Cxcl10 production, which are key in T cell recruitment and may enhance T cell infiltration into the tumor microenvironment.

### PIC promotes Tpex enrichment and Effector CD8⁺ T cell expansion in tumors

To assess whether PIC indirectly modulates T-cell infiltration, tumor-bearing mice were treated with two intratumoral injections of PIC or PBS. Tumors were harvested 20 hours post-treatment and analyzed by FACS, multiplex immunohistochemical consecutive staining on single slide (MICSSS), and scRNA-seq **(Figure 4A)**.

**Figure 4:**
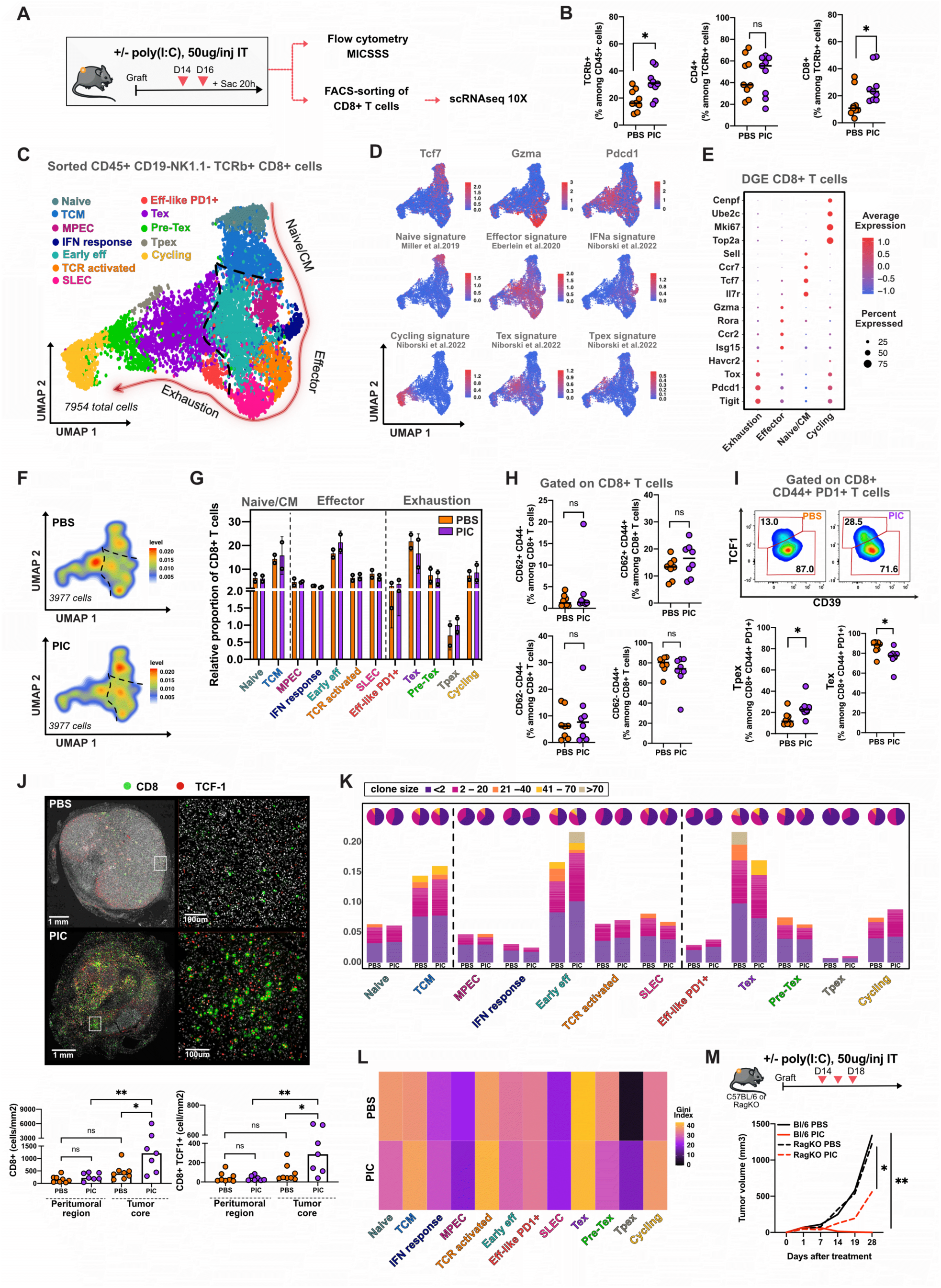
PIC efficacy relies on CD8+ T cells. (**A**) Experimental setup. Tumor-bearing mice were treated intratumorally (i.t) with 50 μg PIC or PBS on day 14 and 16. Tumors were harvested 20h after the last injection and processed for flow cytometry, MICSSS staining, and FACS-sorting of CD8+ T cells for scRNA-seq. **(B)** FACS-based quantification of % of TCRβ⁺ T cells among CD45+ cells, and % of CD4⁺ T cells and CD8⁺ T cells among TCRb+ T cells in PBS- and PIC-treated tumors. Each dot represents an individual tumor. Data shown are representative of two independent experiments, with at least n=8 mice per condition in the experiment presented and n=5 mice per condition in the replicate experiment, both yielding consisting results. Mann-Whitney test. *p<0.05. **(C)** UMAP representation of 7954 tumor-infiltrating CD8⁺ T cells. Defined subsets include naïve/CM (naïve, TCM), Effectors (IFN-response, MPEC, early effectors, TCR-activated cells, and SLEC), Exhaustion (effector-like PD1⁺, Tpex, Tex and Pre-Tex cells) and Cycling. **(D)** Feature plots showing expression levels of gene and transcriptional signatures associated with naïve, effector, IFN response, cycling, and exhausted states (Tpex, Tex). **(E)** Dot plot showing differential gene expression between CD8+ T cell subsets. **(F)** UMAP density plots comparing the distribution of CD8+ T cells populations between PBS and PIC-treated tumors. Top: PBS-treated tumors. Bottom: PIC-treated tumors. 3977 cells per condition. **(G)** Bar graph displaying the relative proportion of naïve/CM, effectors, exhaustion and cycling subsets in PBS and PIC conditions. **(H)** FACS-based quantification of % of naïve (CD62L^+^CD44^-^), TCM (CD62L^+^CD44^+^), effector (CD62L^-^ CD44^+^) and double negative (CD44^-^CD62L⁻) populations among total CD8+ T cells in PBS- and PIC- treated tumors. Each dot represents an individual tumor. Data shown are representative of two independent experiments, with at least n=8 mice per condition in the experiment presented and n=5 mice per condition in the replicate experiment, both yielding consisting results. Mann-Whitney test. *p<0.05. **(I)** FACS-based quantification of % of Tpex (CD44+PD1+TCF1+CD39-) and Tex (CD44+PD1+ TCF1-CD39+) among CD8+ CD44+ PD1+ CD8+ T cells, Top: Representative FACS plots. Bottom: quantification. Each dot represents an individual tumor. Data shown are representative of two independent experiments, with at least n=8 mice per condition in the experiment presented and n=5 mice per condition in the replicate experiment, both yielding consisting results. Mann-Whitney test. *p<0.05. **(J)** Top: Representative MICSSS images of tumor sections from mice treated with PBS (top) or PIC (bottom), stained for DAPI (white), CD8 (green) and TCF1 (red). Zoom-in views (white square) are shown to the right of each image. Scale bars: 1mm (whole tumor), 100um (zoom-in). Bottom: Quantification of total CD8⁺ or CD8+ TCF1⁺ per mm² in tumor core or peritumoral region. PBS n=8 and PIC n=7. Mann-Whitney test. *p<0.05, **p<0.01 **(K)** Distribution of intratumoral CD8+ T cell clones across transcriptional states in two independent PBS- and two PIC-treated tumors. Bar plots show the relative abundance of individual clones within each transcriptional state, stratified by clone size. Corresponding pie charts above each bar represents clonal size. **(L)** Clonal inequality within each transcriptional state measured by Gini index. Higher Gini values indicate dominance of few expanded clones (low diversity), while lower values reflect more even clonal distributions. **(M)** C57Bl/6 or Rag-KO tumor-bearing mice were treated intratumorally with three doses of PIC (50ug/injection) or PBS (control) every other day, starting when tumor volume was between 30-10mm^3^. Top, schematic representation of the experimental workflow. Bottom, tumor growth curves in wild-type (Bl/6) and Rag-KO mice treated with PBS or PIC. Data shown are representative of two independent experiments, with at least n=5 mice per condition in the experiment presented and n=5 mice per condition in the replicate experiment, both yielding consisting results. Mann-Whitney test. *p<0.05, **p<0.01.

Flow cytometry revealed a significant increase in TCRβ⁺ cells in PIC-treated tumors, primarily driven by CD8⁺ T cell levels, while CD4⁺ T cells remained relatively stable **(Figure 4B)**. To further characterize the phenotype of infiltrating CD8⁺ T cells, scRNA-seq was performed on sorted CD45⁺CD19⁻NK1.1⁻TCRβ⁺CD8⁺ T cells from two control and two PIC-treated mice. UMAP projection of the 7,954 resulting cells identified 12 transcriptionally distinct clusters, grouped into four major categories: Naïve/central memory (Naïve/CM), Effector, Exhausted, and Cycling populations **(Figure 4C, S4A–C; Table S5)**.

The Naïve/CM group (including naïve and central memory (TCM) cells) was characterized by expression of *Sell, Il7r, Ccr7*, and *Tcf7*, with low levels of *Gzma* and *Pdcd1*. Effector clusters (IFN-responsive cells, memory precursor effector cells (MPEC), early effectors, TCR-activated cells, and short-lived effector cells (SLEC)) expressed *Gzma*, *Fasl, Ccl5, Rora*, and *Isg15*. Exhausted clusters (effector-like PD1⁺, progenitor exhausted (Tpex), terminal exhausted (Tex) and Pre-Tex cells), were enriched for *Tox, Pdcd1*, and *Tigit*. Cycling cells expressed proliferation markers such as *Cenpf*, *Mki67, Top2a*, and *Ube2c* **(Figures 4D–E, S4C)**. Density plots revealed increased presence of Tpex, early effector, and TCM CD8+ T cell subsets in PIC-treated tumors, along with a notable reduction in Tex cells **(Figure 4F)**. Quantification of relative proportions in the scRNAseq dataset, as well as independent FACS analysis further confirmed a selective enrichment of Tpex cells (PD1⁺CD44⁺TCF1⁺CD39⁻) and a concomitant decrease in Tex cells (PD1⁺CD44⁺TCF1⁻CD39⁺), while the TCM (CD62L+ CD44+) population remained relatively stable **(Figures 4G–I, S4D)**. MICSSS analysis supported these observations, showing increased intratumoral density of CD8⁺TCF1⁺ T cells specifically within the tumor core in PIC-treated mice **(Figure 4J)**.

To determine whether the observed enrichment of Tpex cells stemmed from local expansion or peripheral recruitment, we analyzed the TCR repertoire of intratumoral CD8⁺ T cells. A total of 7854 cells, corresponding to 1653 PBS-clonotypes and 1582 PIC-clonotypes were recovered **(Table S4)**. We visualized the distribution of individual clones based on their abundance using pie charts, and represented the absolute number of cells per clone with bar plots **(Figure 4K)**. Additionally, we assessed clonal expansion within each group by calculating the Gini index, which quantifies the degree of inequality in clonal distribution. High values reflecting dominance of few expanded clones, whereas lower values indicate a more polyclonal repertoire **(Figure 4L)**. In PBS-treated tumors, the highest clonal dominance was observed among Tex cells, consistent with chronic antigen stimulation and progressive clonal expansion leading to exhaustion. In contrast, Tpex showed minimal clonal expansion, suggesting recent recruitment into the tumor and retention of a less differentiated, stem-like state. Upon PIC administration, the most striking changes in clonal dynamics were found among early effector T cells, which displayed marked expansion and included some of the largest clones, likely reflecting the activation of newly recruited, tumor-reactive T cells. Concomitantly, we observed a reduction in the size of the most dominant Tex clones, potentially reflecting their functional elimination due to terminal exhaustion and inability to persist or compete in the reshaped immune landscape **(Figure 4K-L)**. To functionally assess the contribution of T cells to PIC-mediated tumor control, we compared tumor growth kinetics in wild-type (WT) and Rag-deficient mice (Rag-KO). PIC treatment significantly delayed tumor progression in WT mice and, to a lesser extent, in Rag-KO animals, indicating that while T cells are central mediators of the anti-tumor response, additional immune components likely contribute to PIC efficacy (**Figure 4M, S4E).**

Altogether, these results suggest that PIC treatment reprograms the intratumoral CD8⁺ T cell compartment, recruiting lymph node-derived Tpex cells and driving the local expansion of early effector populations, ultimately contributing to tumor control through both T cell-dependent and T-cell independent mechanisms.

### PIC-Induced iNOS⁺ Macrophages Localize Peritumorally and Promote CD8⁺ T Cell Infiltration

Building on our spatial analysis of CD8⁺ T cells, we next investigated the localization and phenotypic characteristics of macrophage populations with potential anti-tumor activity, to determine whether their spatial distribution was associated with immune cell infiltration.

Tumor-bearing mice were treated with a single dose of PIC, and tumors were collected 16 hours post-treatment for analysis by immunofluorescence, MICSSS, and FACS **(Figure 5A)**. To map the spatial distribution of macrophages with potential anti-tumor activity, tumor sections were stained for DAPI, F4/80, and iNOS, the latter serving as a classical marker of inflammatory, anti-tumor macrophage activation^39–41^. High-resolution immunofluorescence imaging **(Figure 5B)** and MICSSS analysis **(Figure S5A)** revealed a striking accumulation of iNOS⁺ macrophages in the peritumoral region, detected exclusively in tumors from PIC-treated mice. This increase in iNOS⁺ macrophages was confirmed by flow cytometry, which showed that approximately 5% of TAMs expressed iNOS following treatment. In contrast, the frequency of CD163⁺ macrophages, a population typically associated with pro-tumoral, immunosuppressive phenotypes- remained stable **(Figure 5C)**. As a note, MICSSS analysis revealed that macrophages represented nearly 80% of the total iNOS-expressing cells, suggesting that the remaining 20% may correspond to other iNOS+ cell within the tumor microenvironment **(Figure S5A)**.

**Figure 5:**
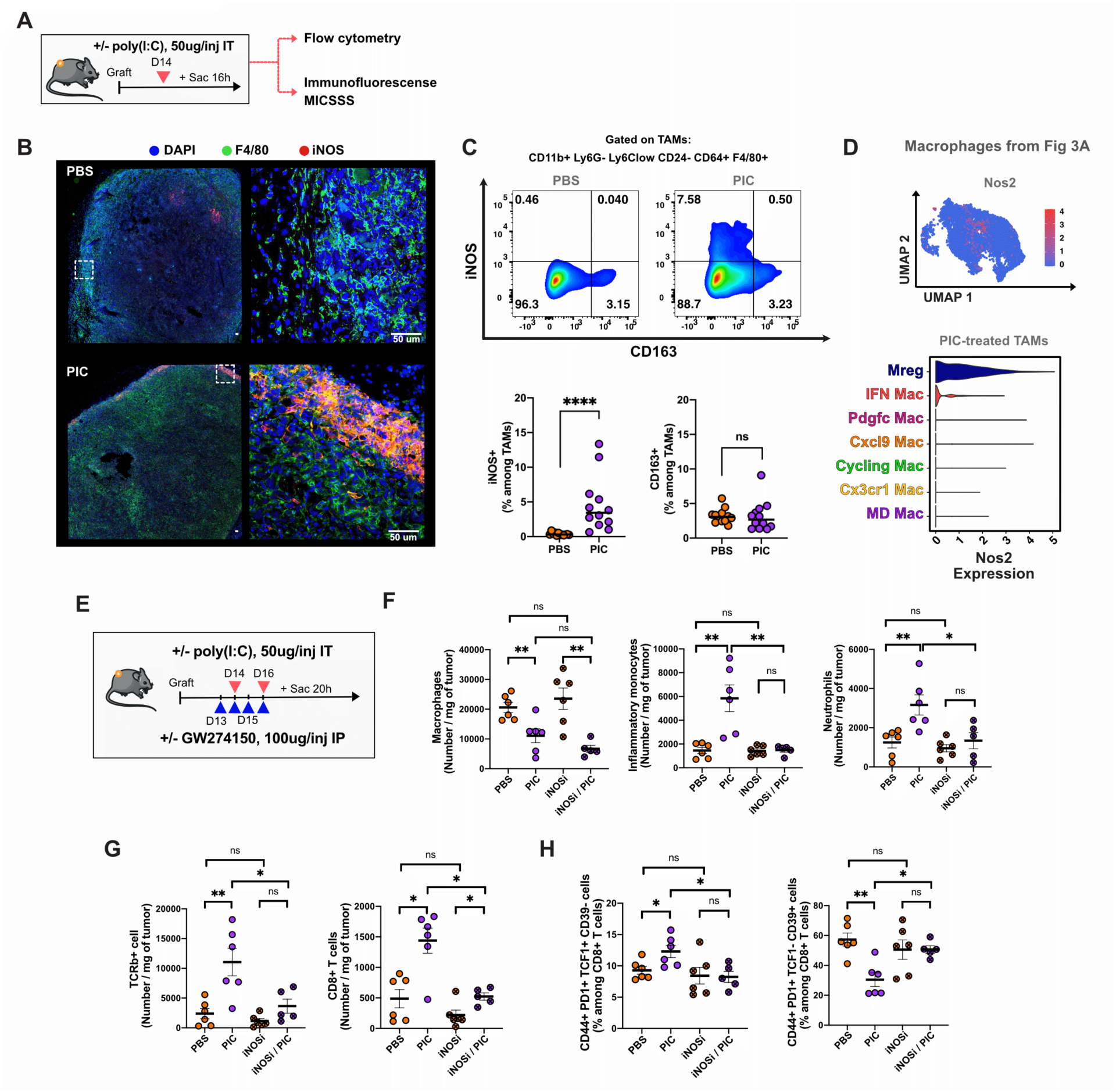
PIC-induced iNOS in peritumoral TAMs is associated to CD8+ Tpex infiltration. **(A)** Schematic representation of the experimental workflow. RT-bearing mice received a single intratumoral (IT) injection of PIC, and tumors were collected 16 hours later for analysis by FACS, immunofluorescence and MICSSS. **(B)** Immunofluorescence images of tumor sections stained for DAPI (blue), F4/80 (green) and iNOS (red), with zoom-in panels (white square) at right. Top: PBS-treated tumors, Bottom: PIC-treated tumors. Scale bar: 50μm. **(C)** FACS quantification of iNOS+ and CD163+ cells on TAMs. Top: Representative FACS plots. Bottom: Percentage among TAMs. Pooled data from two independent experiments including 7 and 5 mice per group, respectively ( total N=12 mice per condition). Mann-Whitney test. ****p<0.0001. **(D)** Nos2 expression in PIC-treated TAMS (in UMAP from figure 3A). Top: Feature plot showing Nos2 expression. Bottom: Violin plots showing Nos2 expression among TAMs clusters. Visualization was done using ALRA assay. **(E)** Schematic representation of the experimental workflow. Tumor-bearing mice were treated intraperitoneally with four doses of GW274150 (daily, 100ug/injection), two intratumoral injections of PIC (every other day, 50ug/injection), with the combination, or PBS as control, starting when tumor volume was between 30-10mm^3^. **(F)** FACS-based quantification of macrophages, inflammatory monocytes and neutrophils in each condition. Graphs show the number of cells per mg of tumor. Mann-Whitney test. *p<0.05, **p<0.01. **(G)** FACS-based quantification of TCRb+ and CD8+ T cells in each condition. Graph represent the number of cells per mg of tumor. Mann-Whitney test. *p<0.05, **p<0.01. **(H)** Frequency of Tpex (TCF1+CD39-) and Tex (TCF1-CD39+) among PD1+CD44+ cells. Mann-Whitney test. *p<0.05, **p<0.01.

To identify the specific macrophage subsets responsible for iNOS expression, we leveraged our previously generated scRNA-seq dataset. iNOS expression was predominantly observed in the Mreg cluster, and to a lesser extent in IFN-responsive macrophages **(Figure 5D)**. Notably, Mregs emerged as the predominant source of heme oxygenase 1 (Hmox1) among all monocyte/macrophage populations, an enzyme that degrades heme to suppress T cell proliferation and promote T cell exhaustion **(Figure S5B)**. Spatial mapping of Hmox1 by MICSSS confirmed that Mregs are enriched in the peritumoral region and capable of expressing iNOS **(Figure S5C)**. Together with their sustained expression of SPP1 post-PIC treatment **(Figure 3H)**, these results indicated that Mreg macrophages maintain their regulatory identity while acquiring inflammatory features such as iNOS expression, suggesting a transitional activation state within this population.

Previous studies have suggested that peritumoral macrophages can regulate immune cell trafficking into tumors^42–44^. Based on this, we hypothesized that iNOS activity could contribute to the recruitment of immune cells in response to PIC treatment. To test this, we pharmacologically blocked iNOS activity using GW274150, a competitive inhibitor of L-arginine that prevents its enzymatic conversion to nitric oxide and citrulline. Mice were treated daily with GW274150, starting one day before PIC administration, and tumors were harvested 20 hours after the second PIC dose **(Figure 5E)**. While iNOS inhibition did not affect the total number of intratumoral macrophages, it significantly reduced the recruitment of inflammatory monocytes, neutrophils, and CD8⁺ T cells **(Figure 5F-G)**. Furthermore, iNOS blockade prevented the enrichment of Tpex cells and abolished the reduction in terminally exhausted (Tex) cells observed following PIC treatment **(Figure 5H)**. Of note, iNOS inhibition also tended to decrease tumor control following PIC treatment **(Figure S5D).** These findings indicate that iNOS activity is required for the recruitment and phenotypic remodeling of CD8⁺ T cell subsets within the tumor.

### PIC synergizes with anti-PD-1 to promote durable tumor control

Building on our observation that PIC reshapes the myeloid compartment and promotes the accumulation of Tpex cells, a CD8⁺ T cell subset associated with responsiveness to PD-1 blockade^45^, we tested whether combining PIC with anti–PD-1 could enhance therapeutic efficacy in vivo.

Rhabdoid tumor–bearing mice were treated with three intratumoral injections of PIC, six intraperitoneal doses of anti-PD-1, or the combination of both, following the schedule depicted in **Figure 6A**. Both monotherapies delayed tumor progression, but the combination therapy achieved significantly superior tumor control **(Figure 6B and S6A)**. Survival analysis revealed better outcome across all treatment groups compared to PBS controls, with the combination therapy providing the most pronounced benefit **(Figure 6C)**. Notably, complete tumor rejection was observed in 8 out of 9 mice in the combination group, versus 5 out of 9 in the PIC group, and 3 out of 9 in the anti-PD-1 group.

**Figure 6:**
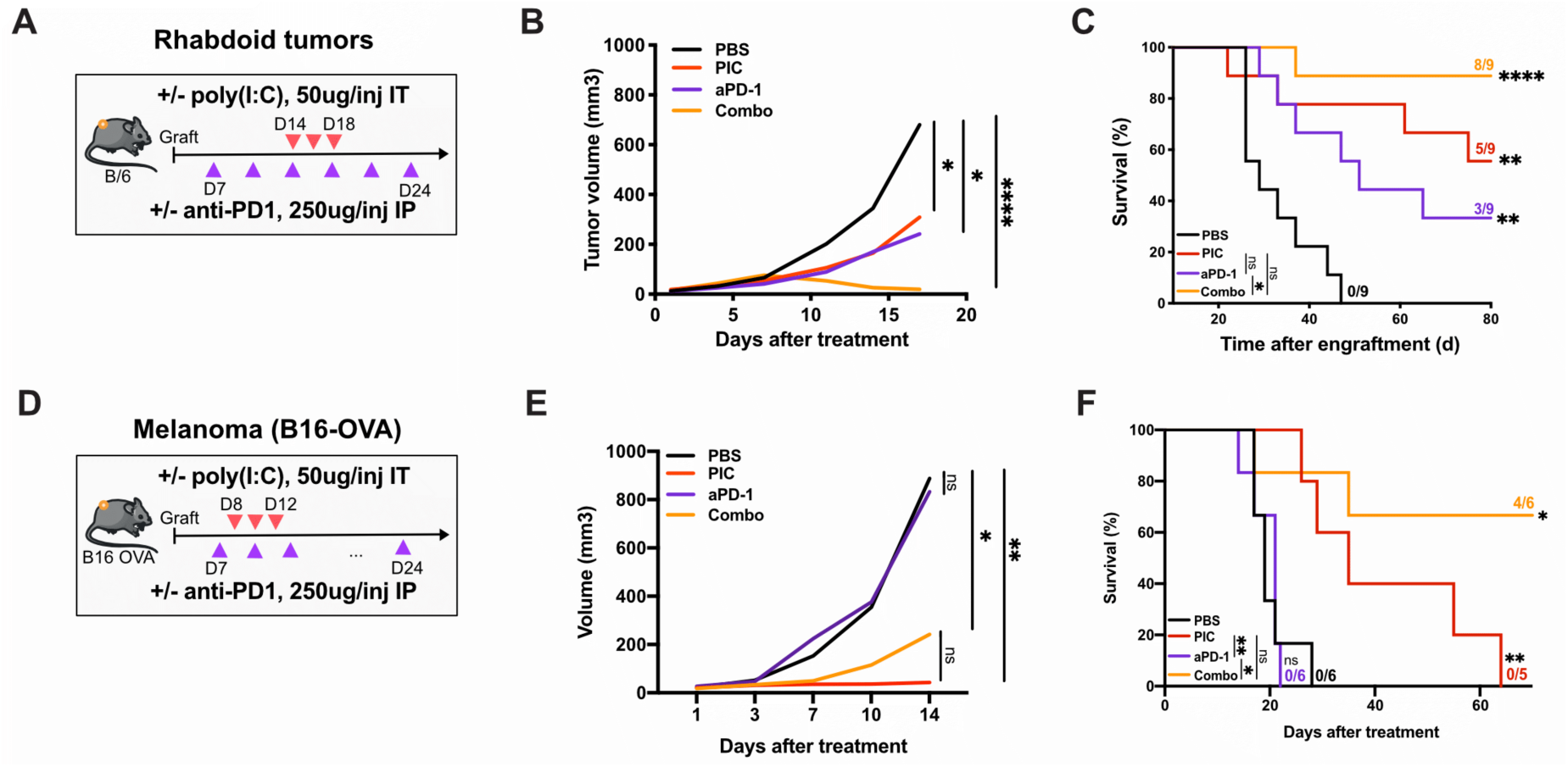
PIC sensitizes tumors to anti-PD1 treatment. **(A)** Schematic of the experimental workflow. Rhabdoid tumor-bearing mice measuring 2–4 mm in diameter (Day 7 post-engraftment) received three intratumoral injections of PIC (50 µg per dose), six intraperitoneal injections of anti–PD-1 antibody (aPD-1, 250 µg per dose) administered twice weekly, or the combination of both treatments (Combo). Control mice received PBS. **(B)** Tumor growth curves for each treatment group in the rhabdoid tumor model. Data represent the mean tumor volume from 9 mice per group in the representative experiment shown. This experiment is representative out of three independent experiments. Statistical significance was assessed using the Mann–Whitney test. p < 0.05, **p < 0.0001 **(C)** Kaplan–Meier survival curves of RT-bearing mice under each treatment condition. Data shown are from the same representative experiment shown in (B). Wilcoxon (log-rank) test; **p** < 0.01, \*\***p** < 0.0001. **(D)** Schematic of the treatment regimen in the B16-OVA melanoma model. Mice bearing subcutaneous B16-OVA tumors (20–100 mm³) received three intratumoral injections of PIC (50 µg/injection), and/or two intraperitoneal injections per week of aPD-1 (250 µg/injection), starting one day prior to the first PIC dose. Control mice received PBS. **(E)** Tumor growth curves for each treatment group in B16-OVA melanoma model. Data represent the mean tumor volume from at least 5 mice per group. Statistical significance was assessed using the Mann–Whitney test. p < 0.05, **p < 0.0001 **(F)** Kaplan–Meier survival curves of B16-OVA-bearing mice under each treatment condition. Data from the same representative experiment shown in (E). Wilcoxon (log-rank) test; ns: not significant, *p* < 0.05, \**p* < 0.01.

To assess whether the synergistic effects of PIC and anti-PD-1 observed in rhabdoid tumors could extend to other poorly immunogenic cancers, we evaluated the combination in the B16-OVA melanoma model, which is classically resistant to immune checkpoint blockade. Treatment was initiated when tumors reached 20–100 mm³, with anti-PD-1 administered one day before PIC **(Figure 6D)**. As expected, anti-PD-1 monotherapy failed to control tumor growth or prolong survival. In contrast, PIC alone elicited a potent antitumor effect, significantly delaying tumor progression and prolonging survival in a subset of mice. However, despite this initial control, relapses were observed following PIC monotherapy **(Figure S6B)**. Notably, combination treatment resulted in both enhanced tumor control and more sustained survival benefits, suggesting that PIC not only exerts direct antitumor activity but also sensitizes otherwise refractory tumors to anti-PD-1 therapy **(Figures 6E–F and S6B)**.

Altogether, these findings demonstrate that the combination of PIC and PD-1 blockade leads to superior and durable tumor control across distinct tumor models, highlighting its strong potential to enhance the efficacy of immune checkpoint therapies.

### Poly(I:C) induces a conserved interferon-driven reprogramming of TAMs across species

To assess whether the transcriptional features of murine PIC-activated TAMs are conserved in human RTs, freshly dissociated tumor samples were stimulated ex vivo for 16h with or without PIC (50 µg/mL), and MoMa were analyzed by scRNAseq **(Figure 7A)**. We examined two representative RT specimens spanning the spectrum of RT subtypes, one primary cerebellar ATRT-SHH and one cerebellar metastasis of an extra-cranial RT (ECRT), thereby allowing direct comparison within a shared brain microenvironment.

**Figure 7.**
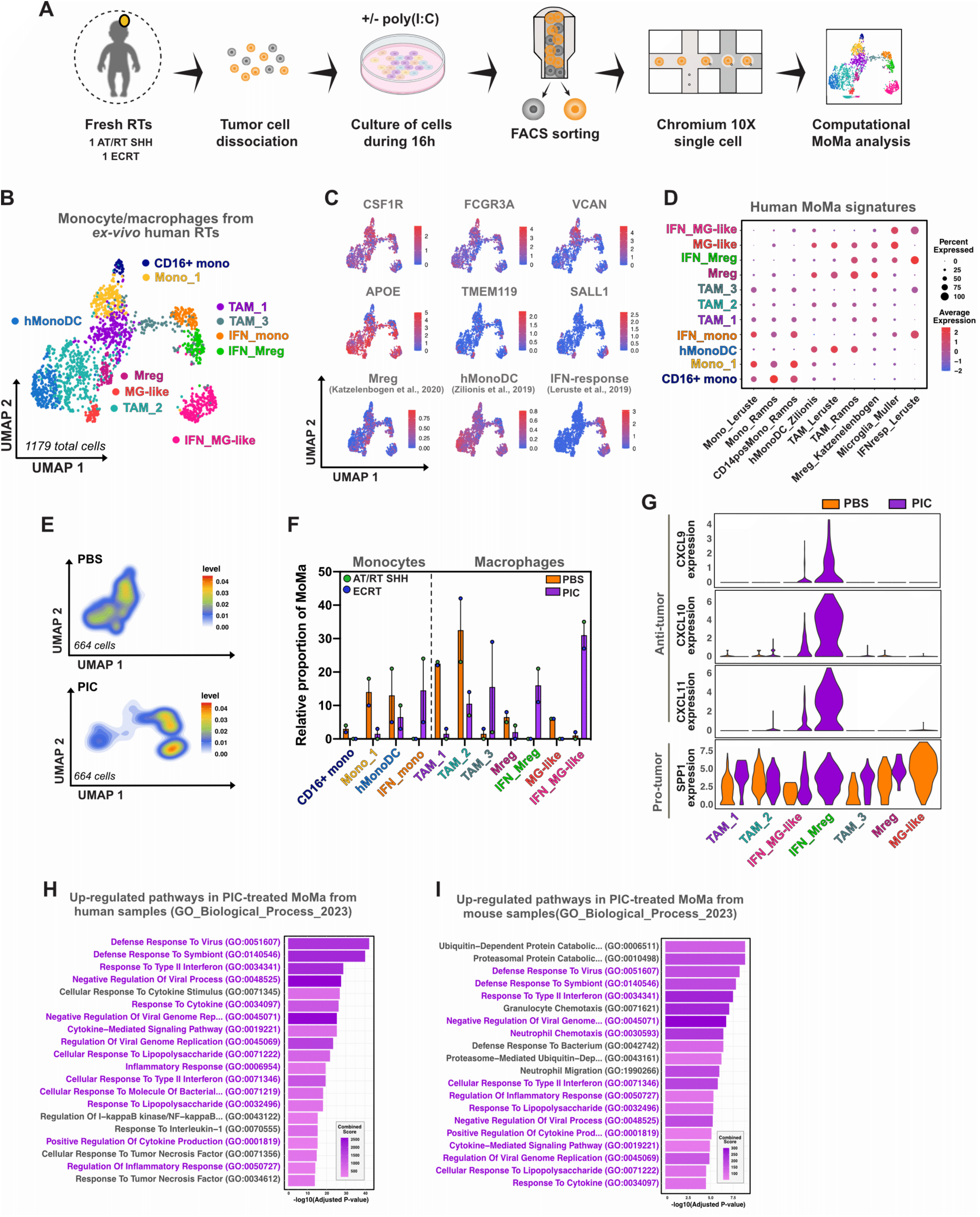
PIC *ex vivo* treatment reprograms human TAMs towards an interferon-driven pro-inflammatory state and reveals conserved responses across species. **(A)** Schematic representation of the experimental workflow. Tumors from two rhabdoid tumor (RT) types (1 ATRT-SHH, 1 ECRT) were dissociated and cultured *ex vivo* for 16 hours with or without PIC (50 µg/mL). Monocyte/macrophage (MoMa) populations were enriched by FACS sorting (CD45+) and subjected to single-cell RNA-sequencing using 10X Genomics Chromium platform. For dowstream analyses, MoMa cells (PTPRC+/CSF1R+/VCAN+/APOE+ and negative for CSF3R, FLT3, CD3E, NCR1) were isolated from the immune compartment. **(B)** UMAP representation of 1,179 MoMa cells, showing refined subclustering. Identified populations include monocytes (Mono_1, CD16⁺ mono, IFN_mono), monocyte-derived dendritic cells (hMonoDC), and diverse tumor-associated macrophage subsets (TAM_1, TAM_2, TAM_3, Mreg, MG-like, IFN_Mreg, IFN_MG-like. Annotation was based on the expression of lineage-defining genes and published signatures. MG was used as abbreviation of microglia. **(C)** Feature plots showing representative genes used to define MoMa subclusters, including canonical monocyte/macrophage markers (CSF1R, FCGR3A), microglial genes (TMEM119, SALL1), and other signatures (IFN-response, Mreg and hMonoDC). **(D)** Dot plot showing the expression of selected subpopulation signatures across MoMa clusters. Dot size represents the percentage of cells within each cluster expressing the corresponding gene, while color intensity reflects the average scaled expression level. Signature were curated from published datasets (Table S6-S7). **(E)** UMAP density plots comparing the distribution of MoMa populations between PBS and PIC- *ex vivo* treated tumors. Top: PBS-treated tumors. Bottom: PIC-treated tumors. n= 664 cells per condition **(F)** Bar plots showing the proportion of monocyte, monocyte-derived, and macrophage clusters across individual tumors, stratified by tumor model (ATRT-SHH and ECRT) and treatment condition (PBS vs. PIC). Data are represented as median. **(G)** Violin plots showing the expression of selected pro-inflammatory and interferon-stimulated chemokines (CXCL9, CXCL10, CXCL1) and the protumoral marker SPP1 across human TAMs subclusters in PBS- and PIC-treated tumors. **(H)** Gene set enrichment analysis (GSEA) of differentially expressed genes (DEGs) in human MoMas isolated from PIC versus PBS-treated tumors. DEGs were identified using thresholds of avg_log2FC > 0.25 and adjusted p-value < 0.05. Enrichment analysis was performed using the GO Biological Process 2023 database. Shared pathways between human and mouse datasets are highlighted in purple. **(I)** Gene set enrichment analysis (GSEA) of differentially expressed genes (DEGs) in mouse MoMas isolated from PIC versus PBS-treated tumors. DEGs were defined as in (H). Enrichment analysis was performed using the GO Biological Process 2023 database. Shared pathways with human datasets are highlighted in purple.

UMAP projection of 1,179 MoMa cells revealed eleven transcriptionally distinct clusters **(Figure 7B)**. Four clusters were classified as monocyte states based on high VCAN and FCGR3A expression and enrichment of established gene signatures: classical monocytes (Mono_1), non-classical monocytes (CD16⁺ mono), monocyte-derived dendritic cells (hMonoDC), and IFN-responding monocytes (IFN_mono). The remaining seven clusters were annotated as macrophage populations according to canonical macrophage/microglial markers (APOE, TMEM119, SALL1) and supported by published subpopulation signatures. These included TAM_1, TAM_2, TAM_3, Mreg, IFN_Mreg, MG-like, and IFN_MG-like **(Figure 7C–7D, S7A, Table S7)**. PIC primarily reshaped the MoMa subpopulations, driving a relative expansion of IFN-programmed clusters, namely IFN_Mreg, IFN_MG-like and IFN_mono, while reducing several monocyte/macrophage subsets, including CD16+ mono, hMonoDC, Mono_1, TAM_1 and TAM2) **(Figure 7E-F)**. This redistribution reflected a shift toward inflammatory, IFN-driving phenotypes. Notably, similar responses were observed in both ATRT-SHH and ECRT samples, despite their distinct molecular subgroup identity **(Figure S7B**). In line with these changes, we examined the expression of IFN-induced chemokines CXCL9, CXCL10, and CXCL11, together with the immunoregulatory macrophage marker SPP1, across TAM clusters. PIC-enriched IFN_Mreg and IFN_MG-like populations displayed strong up-regulation of CXCL9/10/11 while retaining SPP1 expression, consistent with a transitional and plastic phenotype primed for immune activation (**Figure 7G**).

To assess the conservation of PIC-induced transcriptional programs across species, we first identified the differentially expressed genes (DEGs) between PIC- and PBS-treated MoMas in humans and mice (avg_log₂FC > 0.25, adjusted p < 0.05; 538 genes upregulated in humans and 413 in mice). Pathway enrichment analysis based on these DEGs (avg_log₂FC > 0.25, adjusted p < 0.05) revealed extensive overlap between human and murine responses, with 1,598 pathways commonly upregulated (77.3% of mouse and 57.1% of human upregulated pathways) **(Figure S7C, Table S6**). Shared pathways included activation of type I/II IFN responses, antiviral defense, cytokine/chemokine signaling, and leukocyte recruitment, indicating a conserved activation signature following PIC stimulation **(Figure 7 H-I).**

Collectively, these findings demonstrated that PIC reprograms human TAMs into a conserved IFN-driven, pro-inflammatory state, supporting TLR3 activation as a strategy to rewire myeloid cells and sensitize cold tumors to immune attack.

## Discussion

Immune checkpoint inhibitors (ICIs) have achieved unprecedented success in adult malignancies but have shown limited efficacy in pediatric solid tumors, with the notable exception of classic Hodgkin lymphoma^46^. Recent studies suggest that SMARCB1-deficient cancers, such as rhabdoid tumors (RTs), may exhibit increased susceptibility to ICIs through aberrant splicing, reactivation of endogenous retroelements, and sustained interferon signaling^13,46^. However, clinical responses in RTs to PD-1/PD-L1 blockade remain rare and incomplete^15,47^, indicating the presence of additional barriers that restrain therapeutic efficacy.

Single-cell RNA sequencing analysis indicated that pro-tumoral macrophages are the most abundant immune population in both mouse and human RTs^13^, and that together with dendritic cells (DCs) they express TLR3, a receptor sensing double-stranded RNA (dsRNA) whose activation has been shown to induce IFN-β–dependent tumor regression^31^. Meta-analysis of public scRNAseq datasets across pediatric and adult tumors confirmed similar TLR3 expression patterns, underscoring its translational relevance as a conserved myeloid target for therapeutic reprogramming of the tumor microenvironment.

In this study, we treated RT-bearing mice with poly(I:C) (PIC), a synthetic analog of double-stranded RNA (dsRNA), which elicited a robust type I/II interferon response within the tumor microenvironment and achieved potent control of tumor growth. This antitumor effect was mediated through TLR3, rather than cytosolic RNA sensors such as MDA5 or RIG-I^17,18^, highlighting the functional relevance of endosomal RNA sensing in these tumors. TLR3 activation in tumor-associated macrophages (TAMs) reshaped the myeloid landscape and promoted the recruitment of progenitor-exhausted (Tpex) CD8⁺ T cells. Notably, this reprogramming synergized with PD-1 blockade to drive durable tumor regression, unveiling a promising therapeutic strategy to overcome the intrinsic resistance of RTs to immunotherapy.

To delineate how TLR3 activation reshapes the tumor myeloid compartment, we examined the impact of PIC on tumor-associated macrophages (TAMs), revealing two major changes. First, PIC shifted TAMs state toward an antitumor state, demonstrated by robust induction of IFN-responsive chemokines (Cxcl9, Cxcl10, Ccl5) that recruit T cells, accompanied by a concomitant decrease in the mature protumoral Cx3cr1⁺ TAM subset. Consistent with this program, Cxcl9 expression in TAMs has been correlated with improved prognosis and increased intratumoral T-cell infiltration^38,48^, leading to enhanced tumor control and responsiveness to checkpoint blockade^49,50^. In parallel, PIC broadly repressed Spp1 (osteopontin) across most TAM subsets. Given that Spp1 promotes tumor progression by sustaining immunosuppression, angiogenesis, and invasion^51–53^, its downregulation likely reflects a shift toward a less tumor-supportive macrophage state. Consistent with this, elevated Spp1 expression in TAMs has been associated with poor prognosis^38^. Altogether, our results indicate that PIC reduces the burden of protumoral TAMs while reprogramming remaining macrophages toward antitumor phenotypes, a dual myeloid remodeling that may promote tumor control. The notable exception was the myeloid regulatory (Mreg) subset, which further upregulated Spp1 upon PIC exposure in both human and murine macrophages, consistent with reports describing this population as immunosuppressive^54^.

A second distinctive feature of this remodeling was the induction of spatially confined iNOS expression at the tumor border. F4/80⁺ peritumoral macrophages, particularly Mregs, were the main iNOS producers. iNOS catalyzes the production of nitric oxide from L-arginine under inflammatory conditions driven by IFNs^55–57^. Nitric oxide and related reactive nitrogen species can remodel the extracellular matrix and modulate matrix metalloproteinase activity, providing a plausible mechanism by which peritumoral cells relax stromal constraints and facilitate tumor-infiltration^58–60^. Here, we found that iNOS activity in the peritumoral compartment was required for immune recruitment elicited by PIC. Pharmacological inhibition of iNOS abolished the PIC-induced infiltration of inflammatory monocytes, neutrophils, and CD8⁺ T cells, and impaired tumor control. Together, these results identify peritumoral INOS as an essential mediator of tumor accessibility, converting the border from a restrictive barrier into a permissive interface for immune entry, and suggest its potential as biomarker of productive immune activation and response to immunotherapy

Mregs therefore represent a population of particular interest for further study. Despite Nos2 induction upon PIC, they remain Spp1^hi^, indicating a transitional state rather than full inflammatory conversion and highlighting that iNOS alone is not a reliable marker of antitumoral macrophages. This dual phenotype resembles that described by other group^61^, who reported that a subset of tumor-associated macrophages in arrested tumors co-express nitric oxide and arginase, demonstrating that M2-type macrophages can produce NO without losing their M2 identity. Notably, in that context, nitric oxide production correlated with therapeutically effective CD4⁺ T-cell responses, further supporting a link between macrophage-derived NO and productive antitumor immunity. Such hybrid activation states challenge the classical M1/M2 dichotomy and support the notion of functional plasticity within the tumor border. Consistent with this idea, other study^62^ identified axon-tract–associated microglia that maintain cortical boundary integrity and share a transcriptional signature with Mregs, including *Spp1, Gpnmb, Clec7a, Csf1, and Hmox1*. Given their peritumoral localization, we hypothesize that, in addition to a gatekeeping role controlling cellular access, Mregs may help preserve tumor architecture under growth stress, thereby limiting full phenotypic reprogramming upon PIC. Targeted studies will be required to delineate the precise contribution of this population to immune accessibility and structural remodeling at the tumor edge.

Having established that PIC remodels macrophages, we next investigated whether it also engages dendritic cell pathways that license adaptive immunity. In line with^24^, PIC activated cDC1 and enhanced their migration to draining lymph nodes, consistent with increased T-cell priming. Together with the macrophage changes described above, these findings suggest that PIC promotes T cell recruitment into tumors through indirect mechanisms. We observed a selective enrichment of progenitor-exhausted CD8⁺ T cells (Tpex, PD-1⁺ TCF1⁺ CD39⁻), a subset with proliferative potential and the capacity to generate functional effectors^63^. Clonotype analyses supported preferential recruitment from secondary lymphoid tissues rather than local expansion within tumors, in line with recent studies showing that Tpex cells are maintained in tumor-draining lymph nodes and serve as a continuous source of intratumoral effectors^64,65^. To determine whether lymphocytes were required for PIC-mediated tumor control, we next treated Rag-deficient mice lacking T and B cells. Tumor growth inhibition was largely lost in these animals, demonstrating that the antitumor effects of PIC are primarily T cell–dependent, although the innate mechanisms described above may also contribute. Collectively, these results reveal a coordinated cascade in which TLR3-driven myeloid activation establishes the chemotactic and structural conditions for Tpex recruitment, while subsequent T-cell engagement mediates effective tumor clearance.

The enrichment of Tpex cells, a subset associated with responsiveness to PD-1 blockade^45,63,66^, provided a mechanistic basis for combining PIC with checkpoint inhibition. In both RT and the refractory B16-OVA melanoma model, the combination therapy achieved more durable responses than either monotherapy. While PIC alone delayed tumor growth, relapse occurred and the addition of PD-1 blockade not only prolonged survival but also resulted in complete and lasting tumor rejection in most animals, demonstrating that TLR3 activation can convert otherwise resistant tumors into checkpoint-responsive lesions.

Our translational analyses further showed that human RT TAMs respond to PIC ex vivo with a conserved interferon-driven transcriptional program, recapitulating the broad induction of antiviral, cytokine, and chemokine pathways observed in murine MoMas and confirming the remarkable cross-species conservation of TLR3-mediated myeloid activation. The availability of clinically advanced TLR3 agonists, such as Hiltonol and BO-112, with established safety in adults^22,67,68^, offers a clear path toward clinical translation. These agents are already in trials as monotherapies and in combination with checkpoint inhibitors for adult solid tumors^34,69–71^, providing a rationale to extend evaluation to pediatric settings where immunotherapy has historically underperformed.

In summary, our work defines a conserved, IFN-driven mechanism by which TLR3 activation reprograms the myeloid niche, induces iNOS+ peritumoral macrophages, and restores a functional CD8⁺ T-cell compartment, ultimately overcoming resistance to PD-1 blockade. By combining innate sensing to adaptive immune reinvigoration, PIC emerges as a rational combinatorial partner for checkpoint-based therapies with direct relevance for both pediatric and adult “cold” tumors.

### Limitations of the study

This study has certain limitations. Although the number of human ex vivo samples analyzed was limited to two rhabdoid tumors, one atypical teratoid/rhabdoid tumor (ATRT) and one cerebellar metastasis of an extra-cranial rhabdoid tumor, these experiments represent, to our knowledge, the first single-cell transcriptomic analysis of human rhabdoid tumor samples treated ex vivo with poly(I:C) and provide the first indication of how this treatment may act in human RTs. In addition, while our subcutaneous mouse model reproduces key immune–tumor interactions, it does not fully recapitulate the anatomical context in which ATRTs arise. Future evaluation in intracranial rhabdoid tumor models, once available, will be valuable to further assess therapeutic efficacy. Nevertheless, the comparable results obtained in human *ex vivo* and murine tumors suggest that the pathways activated by poly(I:C) are conserved across tumor sites and species, although validation in larger cohorts and clinical settings will still be required.

## Key resources table

| REAGENT or RESOURCE | SOURCE | IDENTIFIER |
| --- | --- | --- |
| **FACS Antibodies** |  |  |
| Anti-mouse CD16/CD32 | BD Biosciences | Cat#553141,<br>RRID: AB_394656 |
| PerCP-eFluor™ 710 anti-mouse CD163, Clone TNKUPJ | Thermo Fisher Scientific | Cat# 46-1631-82,<br>RRID:AB_2716956 |
| FITC anti-mouse Ly-6G, Clone 1A8 | BD Biosciences | Cat#551460;<br>RRID:AB_394207 |
| AF488 Anti-TCF1/TCF7 antibody, Clone C63D9 | Cell Signaling Technology | Cat#6444,<br>RRID:AB_2797627 |
| PE/Cyanine7 anti-mouse CD19, Clone 6D5 | BioLegend | Cat# 115520,<br>RRID:AB_313655 |
| PE/Cyanine7 anti-mouse NK-1.1, Clone PK136 | BioLegend | Cat# 108714,<br>RRID:AB_389364 |
| PE/Cyanine7 anti-mouse TCR $\beta$ chain, Clone H57-597 | BioLegend | Cat# 109222,<br>RRID:AB_893625 |
| PE-Cyanine7 anti-mouse CD11c Clone N418 | Thermo Fisher Scientific | Cat# 25-0114-82,<br>RRID:AB_469590 |
| PE/Cyanine5 anti-mouse I-A/I-E, Clone M5/114.15.2 | BioLegend | Cat# 107612,<br>RRID:AB_313327 |
| PE-eFluor™ 610 anti-mouse iNOS, Clone CXNFT | Thermo Fisher Scientific | Cat# 61-5920-82,<br>RRID:AB_2574636 |
| PE-Texas Red anti-mouse CD11c, Clone N418 | Thermo Fisher Scientific | Cat# MCD11C17,<br>RRID:AB_10373971 |
| PE anti-mouse CD24 , Clone M1/69 | Thermo Fisher Scientific | Cat# 12-0242-82,<br>RRID:AB_465602 |
| APC/Cyanine7 anti-mouse CD45.2, Clone 104 | BioLegend | Cat# 109824,<br>RRID:AB_830789 |
| APC anti-mouse CD64 (FcγRI), Clone X54-5/7.1 | BioLegend | Cat# 139306,<br>RRID:AB_11219391 |
| BUV737 anti-mouse CD11b, Clone M1/70 | BD Biosciences | Cat# 612800,<br>RRID:AB_2870127 |
| Brilliant Violet 785™ anti-mouse Ly-6C, Clone HK1.4 | BioLegend | Cat# 128041,<br>RRID:AB_2565852 |
| Brilliant Violet 711™ anti-mouse Ly6G, Clone 1A8 | BD Biosciences | Cat# 563979,<br>RRID:AB_2738520 |
| Brilliant Violet 650™ anti-mouse/rat XCR1, Clone ZET | BioLegend | Cat# 148220,<br>RRID:AB_2566410 |
| Brilliant Violet 650™ anti mouse F4/80 , Clone BM8 | BioLegend | Cat# 123133,<br>RRID:AB_2562305 |
| Brilliant Violet 421™ anti-mouse CD24, Clone M1/69 | BioLegend | Cat# 101825,<br>RRID:AB_10901159 |
| FITC anti-mouse CD62L, Clone MEL-14 | BD Biosciences | Cat# 553150,<br>RRID:AB_394665 |
| PE-Cyanine7 anti-mouse CD25, Clone PC61.5 | Thermo Fisher Scientific | Cat# 25-0251-82,<br>RRID:AB_469608 |
| PE/Cyanine5 anti-mouse NK-1.1, Clone PK136 | BioLegend | Cat# 108716,<br>RRID:AB_493590 |
| PE-Texas Red anti-mouse CD8 alpha, Clone 5H10 | Thermo Fisher Scientific | Cat# MCD0817,<br>RRID:AB_10374589 |
| PE anti-mouse Ki-67, Clone B56Ki67 | BD Biosciences | Cat# 556027,<br>RRID:AB_2266296 |
| APC/Cyanine7 anti-mouse TCR $\beta$ chain, Clone H57-597 | BioLegend | Cat# 109220,<br>RRID:AB_893624 |
| Alexa Fluor® 700 anti-mouse CD45.1, Clone A20 | BioLegend | Cat# 110724,<br>RRID:AB_493733 |
| APC anti-mouse CD19 | BioLegend | Cat# 115512,<br>RRID:AB_313647 |
| BUV737 anti-mouse CD4, Clone RM4-5 | BD Biosciences | Cat# 612843,<br>RRID:AB_2870165 |
| Brilliant Violet 785™ anti-mouse CD279, Clone 29F.1A12 | BioLegend | Cat# 135225,<br>RRID:AB_2563680 |
| Brilliant Violet 605™ anti-mouse/human CD44, Clone IM7 | BioLegend | Cat# 103047,<br>RRID:AB_2562451 |
| InVivoMAb Anti-mouse PD-1 | BioXcell | Cat#BE0146,<br>RRID:AB_10949053 |
| **Antibodies for Immunofluorescence** |  |  |
| Rabbit anti-iNOS antibody, clone EPR16635 | Abcam | Cat#ab178945<br>RRID:AB_2861417 |
| Rat anti-Mouse F4/80, clone Cl:A3-1 | Biorad | Cat#MCA497R<br>RRID:AB_323279 |
| Donkey anti-Rabbit IgG (H+L) Highly Cross-adsorbed secondary antibody, Alexa Fluor 546. | Invitrogen | Cat#A10040<br>RRID:AB_2534016 |
| Donkey anti-Rat IgG (H+L) Highly Cross-adsorbed secondary antibody, Alexa Fluor 647. | Invitrogen | Cat#A21447<br>RRID:AB_2535864 |
| **Antibodies for MICSSS** |  |  |
| Rabbit anti-F4/80 XP mAB, clone D2S9R | Cell Signaling Technology | Cat#70076S<br>RRID:AB_2799771 |
| Rabbit anti-iNOS antibody, clone EPR16635 | Abcam | Cat#ab178945<br>RRID:AB_2861417 |
| Rabbit anti-Heme Oxygenase 1 antibody, polyclonal | Abcam | Cat#ab299790<br>RRID:AB_299790 |
| Rabbit anti-TCF1/TCF7 antibody, Clone C63D9 | Cell Signaling Technology | Cat# S2203,<br>RRID:AB_2199302 |
| Rabbit monoclonal anti-CD8, Clone D4W2Z | Cell Signaling Technology | Cat# 98941,<br>RRID:AB_2756376 |
| Dako EnVision+/HRP, Rabbit, HRP. Rabbit | Agilent | Cat# K4003,<br>RRID:AB_2630375 |
| **Biological samples** |  |  |
| ATRT-SHH resection | Necker Enfants Malade Hospital and Curie Institute | N/A |
| ECRT metastasis resection | Necker Enfants Malade Hospital and Curie Institute | N/A |
| **Chemicals, peptides, and recombinant proteins** |  |  |
| Poly(I:C) LMW – TLR3 Agonist | InvivoGen | Tlrl-picw |
| Diphtheria toxin (hDT) | Merck | 322326 |
| GW274150 phosphate | MedChemExpress | HY-12119A |
| Liberase TL | Roche | 05989132001 |
| DNase I | Sigma-Aldrich | 100104159001 |
| Paraformaldehyde 4% | Edimex | 3178-200-19 |
| Sucrose 15%/30% | Fisher Scientific | 10634932 |
| OCT compound | Tissue-Tek | 4583 |
| Triton X-100 | Sigma-Aldrich | T8787 |
| Fluoromount-G ( with or without DAPI) | Invitrogen | 15596276/15586276 |
| Peroxidase Blocking Solution | Agilent Dako | S202386-2 |
| Antigen Retrieval Buffer, pH 6.1 | Agilent Dako | S169984-2 |
| Antigen Retrieval Buffer, pH 9 | Agilent Dako | S236784-2 |
| AEC Substrate ( 3-amino-9-ethylcarbazole) | Abcam | Ab64252 |
| Hematoxylin | Agilent Dako | S330930-2 |
| Aqueous Mounting Medium (glycerol-based) | Agilent Dako | C056330-2 |
| Goat serum/Donkey serum | Jackson ImmunoResearch | 005-000-121/<br>017-000-121 |
| BSA IgG-free | Jackson ImmunoResearch | 001-000-162 |
| **Critical commercial assays** |  |  |
| Aquadead | Invitrogen™ | Cat# L34957 |
| Fixation/Permeabilization | eBioscience, Thermo Fisher Scientific | Cat# 88-8824-00 |
| Proteome Profiler Mouse XL Cytokine Array Kit | R&D System | Cat# ARY028 |
| Single Cell 3' Reagent Kit | 10X Genomics | N/A |
| Single Cell V(D)J Reagent Kit (5') | 10X Genomics | N/A |
| **Deposited data** |  |  |
| Murine rhabdoid tumor scRNA-seq and TCR-seq data | This paper | To be deposited at GEO upon publication |
| Human rhabdoid tumor ex vivo scRNA-seq data | This paper | To be deposited at GEO upon publication |
| Human rhabdoid tumor ex vivo scRNA-seq data (Fitte-Duval et al., in preparation) | GEO | To be deposited at GEO upon publication |
| Public human rhabdoid tumor dataset (Leruste et al., 2019) | GEO | Accession: GSE140819 |
| Public human high-grade glioma RNA-seq data (DeSisto et al., 2024) | OpenScPCA, GEO | OpenScPCA code: SCPCP000001, Accession: GSE231860 |
| Public human osteosarcoma RNA-seq data | OpenScPCA | OpenScPCA code: SCPCP000023 |
| Public human neuroblastoma RNA-seq data (Patel et al., 2024; Chapple et al., 2024) | OpenScPCA, GEO | OpenScPCA code: SCPCP000004, Accession: GSE228957 |
| Public human retinoblastoma RNA-seq data (Norrie et al., 2021; Stewart et al., 2017) | OpenScPCA, GEO | OpenScPCA code: SCPCP000011, Accession: GSE174200 |
| Public human tumor-associated myeloid cell atlas (MoMacVerse) scRNA-seq data (Mulder et al., 2021) | GEO | Accession: GSE178209 |
| Public human hepatocellular carcinoma (HCC) scRNA-seq data (Zheng et al., 2017) | GEO | Accession: GSE98638 |
| Public human colorectal adenocarcinoma (COAD) scRNA-seq data (Zhang et al., 2020) | GEO | Accession: GSE146771 |
| Public human pancreatic ductal adenocarcinoma (PDAC) scRNA-seq data (Peng et al., 2019) | GSA | Accession: CRA001160 |
| Public human non-small cell lung cancer (NSCLC) scRNA-seq data (Kim et al., 2020) | GEO | Accession: GSE131907 |
| Public human luminal and triple-negative breast cancer (LBC and TNBC) scRNA-seq data (Azizi et al., 2018) | GEO | Accession:<br>GSE114725 |
| **Experimental models: Cell lines** |  |  |
| B16-OVA (mouse melanoma cell line) | Dr. Eliane Piaggio,<br>Institut Curie | RRID: CVCL_WM77 |
| **Experimental models: Organisms/strains** |  |  |
| *Smarchb1<sup>flox/flox</sup>; Rosa26-CreERT2* mice | Han et al., 2016 | N/A |
| R26S2 syngeneic rhabdoid tumor model | Leruste et al., 2019 | N/A |
| *C57BL/6J* mice | Charles River<br>Laboratories | RRID:IMSR_CRL:027 |
| *CD64-hDTR* | (Baranska et al.,<br>2018) | N/A |
| *TLR3<sup>-/-</sup>* | (Varthaman et al.,<br>2016) | N/A |
| *CARDIF/MAVS<sup>-/-</sup> (C57BL/6NCya-Mavs<sup>em1/Cya</sup>)* | Provided by H.<br>Saklani (Institut<br>Pasteur) | N/A |
| *Rag2<sup>-/-</sup> (B6 Rag2<sup>-/-</sup> KP)* | Alonso et al., 2018 | N/A |
| **Software and algorithms** |  |  |
| Cell Ranger (v6.0.0) | 10x Genomics | https://www.10xgenomics.com/ |
| R (v4.4.1) | R Core Team | https://www.r-project.org/ |
| AnnotationDbi (v1.66.0) | Bioconductor | 10.18129/B9.bioc.AnnotationDbi |
| clustree (v0.5.1) | CRAN | 10.32614/CRAN.package.clustree |
| ComplexHeatmap (v2.20.0) | Bioconductor | 10.18129/B9.bioc.ComplexHeatmap |
| cowplot (v1.1.3) | CRAN | 10.32614/CRAN.package.cowplot |
| dittoSeq (v1.16.0) | Bioconductor | 10.18129/B9.bioc.dittoSeq |
| dplyr (v1.1.4) | CRAN | 10.32614/CRAN.package.dplyr |
| enrichR (v3.4) | Ma'ayan Laboratory | https://maayanlab.cloud/Enrichr/ |
| EnhancedVolcano (v1.18.0) | Bioconductor | https://bioconductor.org/packages/EnhancedVolcano/ |
| ggplot2 (v3.5.1) | CRAN | 10.32614/CRAN.package.ggplot2 |
| ggpubr (v0.6.0) | CRAN | 10.32614/CRAN.package.ggpubr |
| monocle3 (v1.3.7) | Trapnell Lab | http://cole-trapnell-lab.github.io/monocle3/ |
| nichenetr (v2.2.0) | Saeys Lab | https://github.com/saeyslab/nichenetr |
| openxlsx (v4.2.8) | CRAN | 10.32614/CRAN.package.openxlsx |
| org.Hs.eg.db (v3.19.1) | Bioconductor | 10.18129/B9.bioc.org.Hs.eg.db |
| patchwork (v1.3.0) | CRAN | 10.32614/CRAN.package.patchwork |
| pheatmap (v1.0.13) | CRAN | 10.32614/CRAN.package.pheatmap |
| reshape2 (v1.4.4) | CRAN | 10.32614/CRAN.package.reshape2 |
| sctransform (v0.4.1) | Satija Lab | 10.32614/CRAN.package.sctransform |
| Seurat (v4.3.0) | Satija Lab | https://satijalab.org/seurat/ |
| SeuratData (v0.2.2) | Satija Lab | https://satijalab.org/seurat-data/ |
| SeuratWrappers (v0.3.1) | Satija Lab | https://github.com/satijalab/seurat-wrappers |
| SingleCellExperiment (v1.26.0) | Bioconductor | 10.18129/B9.bioc.SingleCellExperiment |
| stringr (v1.5.1) | CRAN | 10.32614/CRAN.package.stringr |
| zellkonverter (v1.14.1) | Bioconductor | 10.18129/B9.bioc.zellkonverter |
| BioRender | BioRender.com | https://biorender.com/ |
| FlowJo (v10.8.1) | BD Biosciences | https://www.flowjo.com/ |
| GraphPad Prism 8.3.1 | GraphPad Software | https://www.graphpad.com/ |
| QuPath(v0.5.1) | Bankhead et al., 2017 | https://qupath.github.io/ |
| ImageJ / FIJI (v2.9.0) | NIH | https://imagej.net/Fiji |
| MICSSS | Giraldo et al., 2018 | DOI: 10.1038/s41596-018-0080-y |
| Affinity Designer (v1.10 .6) | Windows | https://store.serif.com/en-us/update/windows/designer/1/ |

### EXPERIMENTAL MODEL AND STUDY PARTICIPANT DETAILS

#### Human tumor samples

Two freshly resected human rhabdoid tumor (RT) specimens, comprising one atypical teratoid/rhabdoid tumor (ATRT) and one cerebellar metastasis of an extra-cranial RT (ECRT), were collected after obtaining written informed consent from the patients’ parents. All samples were anonymized prior to processing, and relevant clinical information is summarized in **Table S1**. The study protocol, including the consent procedure, was approved by the Institutional Review Boards of the Curie Institute and Necker Hospital for Sick Children (Paris, France; Code projet APHP200122 / N° IDRCB: 2021-A00795-36).

#### Mice

A complete list of mouse strains used in this study is provided in the Key Resources Table. Healthy female C57BL/6J mice were obtained from Charles River Laboratories, housed in a non-barrier facility, and enrolled in experiments at 6–10 weeks of age. Spontaneous tumors were generated using the conditional Smarcb1flox/flox;Rosa26-CreERT2 model as previously described^13,30^, in which biallelic deletion of Smarcb1 was induced by tamoxifen injection at embryonic day (E) 6–7. Syngeneic R26S2 murine RT model was derived from this system and propagated by serial subcutaneous engraftment in the flank every 4–5 weeks. CD64-hDTR mice^72^ were provided by S. Henri (CIPHE, Marseille) via P. Benaroch (Institut Curie) and subsequently crossed to C57BL/6J mice from The Jackson Laboratory. TLR3-KO mice^73^ were generously provided by Monique Lafon (Institut Pasteur) via P. Benaroch (Institut Curie). CARDIF/MAVS knockout mice^74^ (C57BL/6NCya-Mavs^em1^/Cya) were kindly provided by H. Saklani (Institut Pasteur). Rag2^−/−^ mice were obtained from the Bl/6 Rag2^−/−^ KP colony^75^. Experiments using these collaborator-provided models were performed on 6- to 14-week-old male or female mice. Mice were co-housed under standardized conditions with a 12 h light/12 h dark cycle, ambient temperature maintained between 20 and 24 °C, and an average relative humidity of 40–70%. Ethical endpoints were applied for tumor-bearing animals, including a maximal permissible tumor volume of 2000 mm³, >10% body weight loss, or signs of impaired mobility, feeding difficulties, or cachexia. Animal care and use for this study were performed in accordance with the recommendations of the European Community (2010/63/UE) for the care and use of laboratory animals. Experimental procedures were specifically approved by the ethics committee of the Institut Curie CEEA-IC #118 (Authorization APAFIS#16352+201807316073586v2 and #44962-2023092613368865 v1 given by National Authority) in compliance with the international guidelines.

## METHOD DETAILS

### Mice tissue dissociation

Tumors were harvested and digested in 2 mL of CO₂-independent medium (Gibco) supplemented with 0.1 mg/mL DNase I and 0.1 mg/mL Liberase TL (Roche) at 37 °C, using C tubes and a GentleMACS Dissociator (Miltenyi Biotec) with the pre-set program “37C_m_TDK1” (41 min at 37 °C under continuous agitation). Collected lymph nodes were mechanically dissociated prior to enzymatic digestion for 45 min at 37 °C in 2 mL of CO₂-independent medium containing 0.1 mg/mL DNase I and 0.1 mg/mL Liberase TL. Enzymatic reactions were stopped by adding MACS buffer (0.5% BSA and 2 mM EDTA in PBS), and cell suspensions were filtered through a 100 µm cell strainer (Falcon). Red blood cells were removed by treatment with RBC lysis buffer (NH4 Cl 155mM, NaHCO_3_ 10mM, EDTA 9,12mM). The resulting single-cell suspensions were washed, resuspended in MACS buffer, and the number of viable cells was determined using acridine orange staining with an automated cell counter (LUNA-FX7, Logos Biosystems).

### FACS Analysis and Cell Sorting on Mouse RT Models

For flow cytometry analysis, up to 4 × 10⁶ cells were resuspended in 100 µL of MACS buffer per sample. Cells were incubated at 4 °C for 10 min with mouse Fc Block (anti-CD16/CD32, clone 2.4G2) to prevent nonspecific antibody binding, washed with MACS buffer, and subsequently stained at 4 °C for 30 min with the Live/Dead Fixable Aqua Dead Cell Stain Kit (Life Technologies) to discriminate live and dead cells. The antibody panel used for cell surface labeling (see Key Resources Table) varied depending on the experimental setting and tissue analyzed but were always incubated at 4 °C. For intracellular staining, cells were permeabilized and fixed using the Fixation/Permeabilization Kit (eBioscience, Thermo Fisher Scientific) according to the manufacturer’s instructions. Data acquisition was performed on an LSR Fortessa cytometer (BD Biosciences), and analyses were carried out using FlowJo software (Tree Star). The gating strategy is illustrated in Figures S2F, S2I and S4D.

For cell sorting, all samples were stained at 4 °C with a specific antibody cocktail tailored to each experimental condition, along with DAPI to exclude dead cells, and subsequently filtered through a 70 µm cell strainer. Cells were sorted at 4 °C using a FACSAria III or a FACSAria Fusion cytometer (BD Biosciences) equipped with a 100 µm nozzle, and collected into serum-precoated Eppendorf tubes containing PBS supplemented with 0.04% BSA. Sorted cells were processed within 2 hours and loaded onto a 10x Genomics Chromium instrument for downstream applications.

### Mice treatment procedure

For the generation of syngeneic RT models, 10–20 mm³ tumor fragments were engrafted subcutaneously into the flank of recipient mice. Starting on day 7 post-engraftment, mice were monitored, and those with measurable tumor size were included in the experiment. For each experiment, the start date of treatment varied because intratumoral injections required tumors to reach an adequate size for accurate administration. On the first day of treatment, groups were stratified based on tumor volume to ensure equivalent mean tumor size across experimental arms. Tumor volume was determined by caliper measurements three times per week using the formula (length × width²)/2. Survival under treatment was assessed until the ethical endpoints described previously.

#### In vivo macrophage depletion

For macrophage depletion experiments, 2- to 5-month-old male and female CD64 wild-type (CD64^WT^) and CD64-hDTR (CD64^hDTR^) tumor-bearing mice with tumor volumes ranging from 30 to 100 mm³ at day 14 were treated intraperitoneally with four doses of human diphtheria toxin (1 µg in 100 µL per injection; hDT, Merck, 322326) administered on days 14, 16, 18, and 20 post-engraftments.

#### Tumor Progression in PIC-Treated Mice

Two-month-old female C57BL/6 tumor-bearing mice with tumor volumes ranging from 30 to 100 mm³ at day 14 post-engraftment were treated intratumorally with three doses of PIC (50 µg in 50 µL per injection; InvivoGen, tlrl-picw) administered on days 14, 16, and 18, or with 50 µL of intratumoral PBS as a control.

For flow cytometry analysis of myeloid populations, tumors were harvested 16 hours after a single dose of PIC administered on day 14 post-engraftment. In some experiments, tumor fragments of equal weight were mechanically dissociated in 500 µL of PBS, and the resulting supernatants were analyzed using the Proteome Profiler Mouse XL Cytokine Array Kit (R&D Systems). For flow cytometry analysis of T cells, tumors were collected 20 hours after the second PIC injection administered on day 16 post-engraftment.

#### Tumor progression in TLR3 KO and MAVs KO mice

For experiments assessing tumor progression in TLR3- and MAVS-deficient mice, 2- to 8-month-old tumor-bearing TLR3 wild-type (TLR3^WT^) and TLR3 knockout (TLR3^KO^) mice, as well as MAVS wild-type (MAVS^WT^) and MAVS knockout (MAVS^KO^) mice, with tumor volumes ranging from 30 to 100 mm³ at day 14 post-engraftment, were treated intratumorally with three doses of PIC (50 µg in 50 µL per injection) administered on days 14, 16, and 18.

#### Tumor progression in Rag-deficient mice

For experiments to functionally assess the contribution of T cells to PIC-mediated tumor control, 2- to 8-month-old tumor-bearing wild-type (WT) and Rag-deficient mice (Rag-KO), with tumor volumes ranging from 30 to 100 mm³ at day 14 post-engraftment, were treated intratumorally with three doses of PIC (50 µg in 50 µL per injection) administered on days 14, 16, and 18.

#### In vivo iNOS blockade

For iNOS blockade Two-month-old female C57BL/6 tumor-bearing mice were treated intraperitoneally with seven daily doses of GW274150 (1 µg in 100 µL per injection) starting the administration at day 13 after engraftment, three intratumoral injections of PIC (50 µg in 50 µL per injection) on days 14,16 and 18, with the combination, or PBS as control. For FACS experiments, tumors were collected after four doses of iNOS inhibitor and two doses of PIC.

#### Tumor progression in mice treated with anti-PD1 and PIC

Syngeneic RT models with tumor diameters between 2 and 4 mm at day 7 post-engraftment were stratified by tumor size to ensure equivalent mean tumor volume across the following four groups: PBS, anti-PD-1, PIC, and combination treatment. Mouse anti-PD-1 antibody (clone RMP1-14; BioXCell) was administered intraperitoneally at a dose of 250 µg per mouse on days 7, 10, 14, 17, 21 and 24 post-engraftment. PIC was administered intratumorally at a dose of 50 µg per mouse on days 14, 16, and 18, and PBS was administered as the control treatment. For the B16-OVA tumor models, 500,000 tumoral cells were resuspended in 100 µL of PBS and subcutaneously injected into the right flank of female mice. Treatments started once tumors reached an initial tumor volume ranging from 20 to 100 mm³, which typically occurred around seven days post-inoculation. The mice were stratified as done in the RT model. Mouse anti-PD-1 antibody (clone RMP1-14; BioXCell) was administered intraperitoneally at a dose of 250 µg per mouse bi-weekly starting one day before than PIC, while PIC was administered intratumorally at a dose of 50 µg per mouse on days 8, 10, and 12. PBS used as control.

### Single-Cell RNA Sequencing

FACS-enriched mouse live cells, mouse myeloid cells (CD45⁺ TCRβ⁻ NK1.1⁻ CD19⁻), and human tumor-infiltrating cells treated ex vivo with PIC were loaded onto a 10X Chromium system (10x Genomics), and libraries were prepared using the Single Cell 3′ Reagent Kit v2 (10x Genomics) according to the manufacturer’s protocol. Library quantification and quality assessment were performed using the dsDNA High Sensitivity Assay Kit (Thermo Fisher Scientific) and a Bioanalyzer 2100 system (Agilent Technologies), and libraries were sequenced on an Illumina HiSeq 2500 platform.

FACS-enriched mouse CD8⁺ T cells (CD45⁺ NK1.1⁻ CD19⁻ TCRβ⁺ CD8⁺) were processed for single-cell TCR sequencing coupled to 5′ gene expression profiling using the Single Cell V(D)J Reagent Kit (10x Genomics) following the manufacturer’s instructions. Briefly, after amplification and purification of barcoded full-length cDNA, V(D)J segments were selectively enriched prior to library construction. 5′ gene expression libraries were sequenced on an Illumina NovaSeq platform, and enriched V(D)J libraries were sequenced on an Illumina MiSeq system. The number of cells loaded and recovered for each experiment is provided in the **Table S1, S2, S3 and S5**.

### Single-cell RNA-seq data analysis

In general, single-cell RNA-seq data were processed using Cell Ranger v6.0.0 (10x Genomics) with the refdata-gex-mm10-2020-A reference genome for mouse datasets and the refdata-gex-GRCh38-2020-A reference genome for human datasets. The resulting gene-barcode UMI count matrices were analyzed in RStudio (v4.0) with Seurat v4.

Quality control metrics and dataset-specific thresholds were applied individually to each sample. Initial filtering was based on the polymodal distribution of cells by log2-transformed percentage of mitochondrial gene content and log2 total UMI counts per cell, to exclude dead or uninformative cells. Specific cutoffs for each dataset are detailed in the Supplementary Tables associated with each figure. For CD8+ T cell analyses, both mitochondrial and gene count thresholds were applied simultaneously. In all other datasets, no initial gene count threshold was applied prior to the identification of neutrophil populations, given their inherently low transcriptional complexity.

Normalization was performed using Seurat’s SCTransform function^76,77^. Data integration was conducted with the anchor-based workflow implemented in Seurat^78^. This approach identifies pairs of cells in similar biological states (“anchors”) across different samples (e.g., individuals or experimental conditions) to harmonize gene expression profiles. Integration was performed using the FindIntegrationAnchors() and IntegrateData() functions. No additional batch correction methods were applied.

To reduce technical noise and dimensionality, principal component analysis (PCA) was computed on the scaled expression matrix using the RunPCA() function, and the most informative principal components were retained for downstream analyses. For CD8+ T cell data, TCR genes were excluded from dimensionality reduction and clustering to avoid biases introduced by clonal expansion.

Graph-based clustering was performed using the Louvain algorithm via FindClusters() functions, and visualized using Uniform Manifold Approximation and Projection (UMAP) implemented in the RunUMAP() function. Clustering stability across resolutions was assessed with Clustree v0.5.1.

Canonical lineage markers were used to identify contaminating cell populations, which were removed from the dataset. All downstream analyses, including normalization, integration, clustering, and visualization, were then repeated after contaminant removal.

### Cell type annotation and Downstream Analysis

Cell clusters were annotated based on differential gene expression analysis performed using the FindAllMarkers() function (Wilcoxon rank-sum test, adjusted P-value <0.05, |log2 fold change| >0.25) and the expression of canonical cell population markers. For validation and refinement of cluster identities, curated gene signatures derived from published literature were quantified at the single-cell level using the AddModuleScore() function. Human and mouse signatures used in this study were listed in **Table S7** ^36,79–98^.

<u>For the analysis of druggable targets</u>, non-treated CD45+ cells were clusterized according to main immune populations, and a heatmap representation of the relative mean expression of selected myeloid druggable targets (Cd274 (PD-L1), Csf1r, Pdcd1lg2 (PD-L2), Sirpa, and Tlr3, across different clusters was done using normalized cluster-wise mean expression values.

<u>Trajectory</u> inference of monocyte and macrophage differentiation was performed using Monocle3^99^, designating root nodes corresponding to monocyte clusters in accordance with established lineage relationships^100^.

<u>Pathways analysis</u>. differential gene expression analysis between conditions in this subset was performed (Wilcoxon rank-sum test, adjusted p < 0.05, |log₂FC| > 0.25). According to these, upregulated (PIC) and downregulated (PBS) genes were determined for pathway enrichment analysis, using Enrichr’s API. Enrichment calculations were carried using the Gene Ontology (GO) Biological Process and Molecular Function 2023 databases, the top 10 upregulated and downregulated terms then were visualized using bar plots, ranked by -log10(adjusted p-value).

<u>To analyse and compare lowly expressed genes</u>, we applied the Adaptively-thresholded Low Rank Approximation (ALRA) method^101^. ALRA selectively imputes technical zeros, caused by dropout events, while preserving true biological zeros. This approach enhanced data visualization and allowed us to better compare the expression of genes such as iNOS across our different conditions. The use of ALRA was indicated in the legend of the figure.

### TCR-seq Data processing and Analysis

V(D)J sequences were assembled using Cell Ranger v6.0.0 (10x Genomics) aligned to the refdata-cellranger-vdj-GRCm38-alts-ensembl-7.0.0 reference genome. Clonotypes were defined on the basis of unique CDR3 nucleotide sequences, and only full-length, productive TRA and TRB contigs were retained for downstream analyses. Cells harboring more than one TRB or more than two TRA chains were excluded as potential doublets.

Clonotype expansion was quantified by enumerating the number of cells expressing each clonotype. Data were grouped by clonotype, treatment condition, and annotated T cell subpopulation to assess clonal expansion in response to experimental perturbations. Clonal composition and relative abundance were visualized using bar plots and pie charts to facilitate comparison across conditions.

To estimate clonal diversity, the Gini-TCR Skewing Index was calculated as previously described^102^, ranging from 0 (uniform clonal distribution) to 100 (highly skewed repertoires dominated by few clonotypes). This index was computed separately for each T cell subpopulation and treatment group.

### Metanalysis of TLR3 expression

For the meta-analysis of TLR3 expression in single-cell data, we extracted pediatric tumor single-cell datasets from the Single-cell Pediatric Cancer Atlas Portal of Alex’s Lemonade Stand Foundation. For adult monocyte and macrophage data, we utilized the MoMac-VERSE database from Gustave Roussy’s Florent Ginhoux’s Lab. In both cases, the expression matrices and corresponding metadata were retrieved and processed into Seurat objects for downstream analysis of TLR3 expression. For the pediatric cancer datasets, cluster harmonization was necessary to enable cross-dataset comparisons. Specifically, original clusters with similar cell type annotations (e.g., “Macrophage 1” and “Macrophage 2”) were consolidated into unified clusters (e.g., “Macrophages”). This step was not required for the adult datasets, as they were consistently annotated across samples. Finally, TLR3 expression levels were compared across cell types using dot plots, which facilitated visualization and interpretation of expression patterns.

### Gene set enrichment and cross-species pathways analysis of PIC-induced responses in human and mouse MoMa

To evaluate the conservation of PIC-induced transcriptional programs between human and murine MoMas, differentially expressed genes (DEGs) were identified between PIC- and PBS-treated samples for each species using the Wilcoxon rank-sum test (adjusted p < 0.05, avg_log₂FC > 0.25). A total of 538 and 413 upregulated genes were obtained for human and mouse MoMas, respectively.

Pathway enrichment analyses were then performed separately for each species using the Enrichr API and the Gene Ontology (GO) Biological Process and Molecular Function 2023 databases. Significantly enriched pathways (adjusted p < 0.05) were ranked by –log₁₀(adjusted p-value) and visualized using bar plots. Finally, upregulated pathways from both species were compared to identify conserved responses. Overlapping pathways were quantified and categorized, revealing shared enrichment in type I/II interferon signaling, antiviral defense, cytokine/chemokine signaling, and leukocyte recruitment.

### Immunofluorescence

Tumors were collected in 4% paraformaldehyde (Edimex, 3178-200-19) and incubated for 24 h at 4 °C in the dark. Samples were dehydrated in two successive baths of 15% and 30% sucrose (Fisher Scientific, 10634932), embedded in optimum cutting temperature compound (OCT; Tissue-Tek, 4583), and stored at –20 °C. Tumors were sectioned at 20 µm thickness. Sections were permeabilized using 0.2% Triton X-100 (Sigma, T8787) diluted in PBS and blocked with a solution containing 1:100 Fc Block (BD, 553142), 1:20 goat serum (Jackson ImmunoResearch, 005-000-121) or donkey serum (Jackson ImmunoResearch, 017-000-121) depending on the host species of the secondary antibodies, and 3% IgG-free bovine serum albumin (Jackson ImmunoResearch, 001-000-162) in 0.05% Triton X-100 in PBS. Sections were stained with primary antibodies against F4/80 (Biorad, MCA497R) and iNOS (Abcam, ab178945). Corresponding secondary antibodies were donkey anti-goat AF647 (Invitrogen, A21202; 1:200) and donkey anti-rabbit AF546 (Invitrogen, A10040; 1:100). Slides were mounted using Fluoromount-G™ with DAPI (Invitrogen, 15596276). Images were performed on an inverted two-photon laser-scanning confocal microscope (Leica DMi8, SP8 scanning head unit) equipped with HC PL APO CS2 40x/1.30 OIL objective. Acquisition was performed in resonant mode with a Z-stack thickness of 10 µm and a Z-step size of 2 µm. Tiling arrays covering the entire tumor were acquired. Image stitching was performed using LAS X software (Leica Microsystems).

### Multiplex immunohistochemistry

Tumors were collected in 4% paraformaldehyde (Edimex, 3178-200-19), incubated for 24 h in the dark and then prepared by the platform of Experimental pathology of Institut Curie. Multiplexed immunohistochemistry (IHC) was performed according to the protocol developed by Remark et al. (2016), with minor adjustments. Formalin-fixed paraffin-embedded tumor sections (3 µm thickness) were baked for 1 h at 60 °C, followed by deparaffinization and rehydration in successive baths of xylene and descending gradients of ethanol (100%, 90%, 70%, 50%, and deionized water). Heat-induced epitope retrieval was performed in a water bath at 95 °C for 30 min for the first staining cycle (and 15 min for subsequent cycles), using either pH 6.1 (Agilent Dako, S169984-2) or pH 9 (Agilent Dako, S236784-2) retrieval buffers.

Tissues were stained manually. Sections were incubated in peroxidase blocking solution (Agilent Dako, S202386-2) for 10 min, followed by a Fab fragment block for 20 min if the primary antibody was from the same species as antibodies used in previous cycles. A protein block was applied for 10 min prior to incubation with primary antibodies for 1–2 h. Detection was performed using species-specific secondary antibodies conjugated with horseradish peroxidase (anti-rabbit: Agilent Dako, K400311-2; anti-mouse: Agilent Dako, K400111-2; anti-rat: BioTechne, VC005-050). Chromogenic revelation was achieved with 3-amino-9-ethylcarbazole (AEC; Abcam, ab64252). Slides were counterstained with hematoxylin (Agilent Dako, S330930-2) and mounted using a glycerol-based aqueous mounting medium (Agilent Dako, C056330-2). Following each staining and imaging cycle, slides were scanned using a NanoZoomer S360 scanner (Hamamatsu, C13220-01). Coverslips were removed, and the chromogenic stain was stripped by ethanol baths before the next cycle, beginning again with heat-induced epitope retrieval.

### Multiplex immunohistochemistry imaging analysis

Stains acquired from the same sample were deconvolved into pseudo-fluorescence channels using the HALO Indica Labs Deconvolution v1.1.8 module. Images were registered with the HALO Registration module and fused into a single composite overlay in TIFF format. Overlays were cropped and exported as OME-TIFF files for analysis in QuPath software^103^.

The peripheral tumor region was delineated by selecting the first 500 µm from the tumor edge inward. The remaining area was defined as the tumor core. To avoid analytical biases, regions containing tears or artifacts were excluded from analysis. Nuclei detection was performed using the Halo Software (V3.5; Indica Labs) or StarDist plugin^104^. Positive cell detection was carried out using standard QuPath functions. Extracted data were further analyzed using dedicated statistical and image analysis workflows.

## QUANTIFICATION AND STATISTICAL ANALYSIS

The tests used for statistical analyses are described in the legends of each concerned figure and have been performed using GraphPad Prism v8 or R v4.3. Symbols for significance: ns, non-significant; *, <0.05, **, <0.01; ***, <0.001; ****, <0.0001. For each experimental group, n represent the number of subjects within each group.

## Author contributions

Conceptualization, V.M., E.P., F.B., and J.J.W.; Performed Experiments, V.M., S.C.S., L.L.N., A.L., C.S., J.D., J.M., Z.F., L.L.L, Y.G-F., S.F-D., F.M., R.M-O., J.T-B., R.B-B., M.F.P., M.B and S.B.; Formal Data Analysis, V.M., S.C.S., L.L.N., A.L., J.M., L.L.L, W.R., M.V., and Y.M-K.; Resources, K.B., J.H., P.B., C.S.,J.G and H.M.; Assembled Figures (Visualization), V.M.; Data Curation, V.M.; Writing Original Draft, V.M. ; Writing, Review & Editing, V.M., E.P., F.B., J.J.W., P.B., C.S and J.H.; Project Administration, E.P., F.B., and J.J.W.

## Acknowledgements

This work was funded by Fondation pour la Recherche Médicale (FRM, grant number EQU202203014661), the “Agence Nationale de la recherche” (ANR-15-C E13-009), Janssen Horizon, “Ligue nationales contre le Cancer” (ELCK20384), PRTK-19-052 funding from INCa and DGOS, the LabEx DCBIOL (ANR-10-IDEX-0001-02 PSL; ANR-11-LABX-0043); and Center of Clinical Investigation (CIC IGR-Curie 1428). We would like to thank Marabout de Ficelles and SFCE-Enfants Cancer Santé and Recherche clinique, Entrepôts de données et Pharmacologie GHU Paris Centre Université Paris Cité, URC, Paris Kids Cancer, React4kids and the unité de recherche clinique from Necker Hospital, Paris, for supporting all our research for rhabdoid tumors. We also acknowledge La Ligue contre le cancer for funding the PhD programs of Stéphanie Fitte-Duval and Sofia Cavada Silva. We would like to acknowledge the Institut Curie’s CurieCoreTech platforms for their technical and scientific expertises. We thank Virginie Dangles-Marie, Christopher Dez, Mickaël Garcia and all members of animal facility and Coralie Guérin, Léa Guyonnet, Anne-Gaëlle Lafont, Anna Chipont, Annick Viguier, at Curie’s Cytometry platform (CYTPIC). High-throughput sequencing was performed by the ICGex NGS platform of the Institut Curie supported by the grants ANR-10-EQPX-03 (Equipex) and ANR-10-INBS-09-08 (France Génomique Consortium) from the Agence Nationale de la Recherche (“Investissements d’Avenir” program), by the ITMO-Cancer Aviesan (Plan Cancer III) and by the SiRIC-Curie program (SiRIC Grant INCa-DGOS-465 and INCa-DGOSInser12554). Data management, and quality control were performed by the Bioinformatics platform of the Institut Curie. We also thank Sonia Lameiras, Virginie Raynal, Benoit Albaud, and Patricia Legoix from NGS Platform and to the Platform of Experimental Pathology (PATEX), specially to André Nicolas. We also acknowledge Celio Pouponnot, Sara Chabi, Mathieu Maurin, Maeva Veyssiere and Owen Hoare for their helpful discussions. Finally, we thank the Lemonade Stand Foundation and Florent Ginhoux’s laboratory for providing open access to their databases, which made our meta-analyses possible.

## Declaration of generative AI and AI-assisted technologies in the writing process

During the preparation of this work, the authors used ChatGPT (OpenAI) in order to assist in improving the readability and clarity of the manuscript. After using this tool or service, the authors reviewed and edited the content as needed and take full responsibility for the content of the publication.

---

**Supplementary Figure 1.**
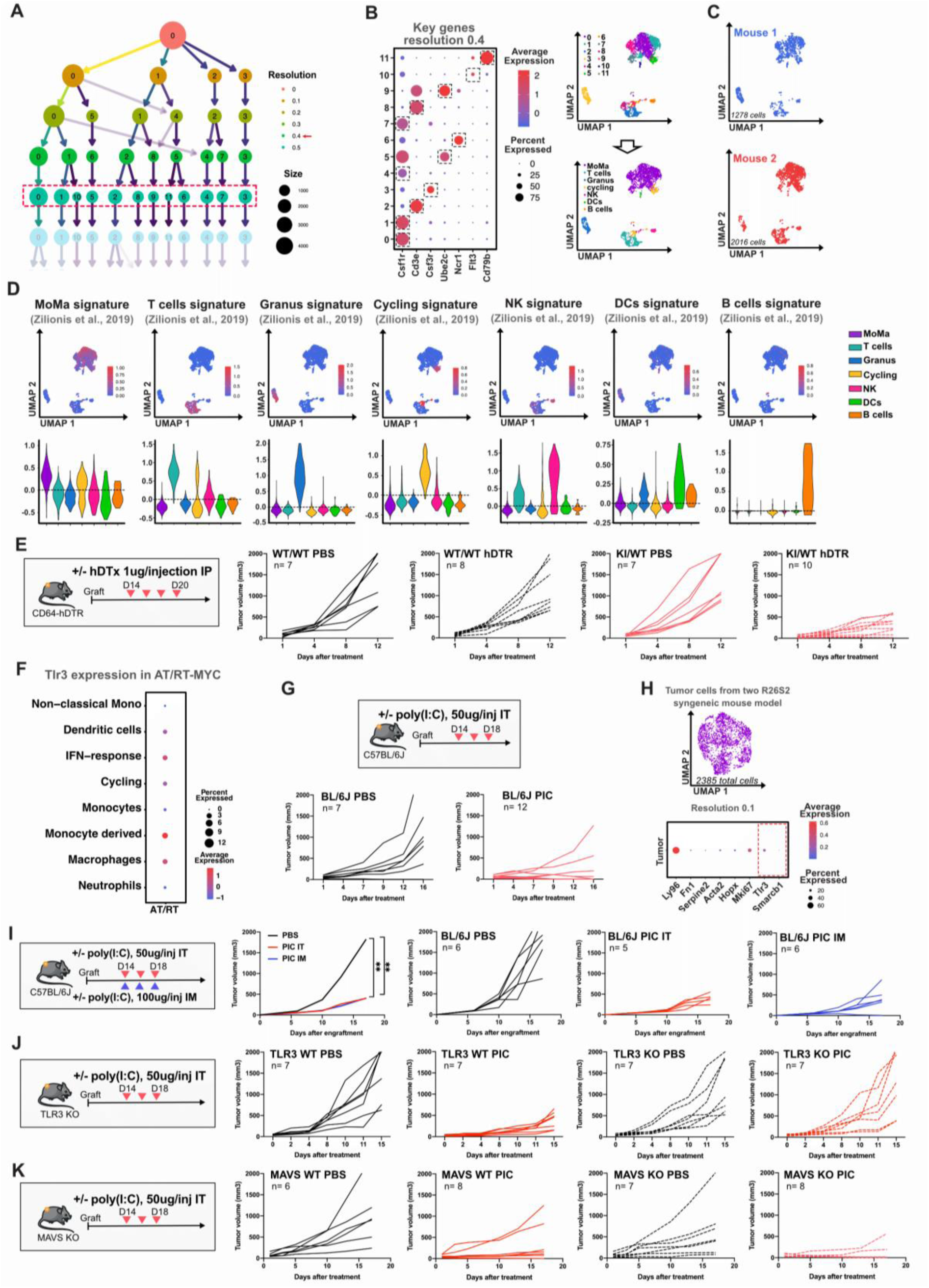
**A)** Clustree highlighting the selected resolution of 0.4 (marked in red dotted lines square), which identified eleven distinct clusters. The dotted rectangle highlights the selected resolution. **B)** Left, Dot plot illustrating the scaled expression of key genes for each subpopulation. Dot size indicates the percentage of cells in each cluster expressing the corresponding gene. Cells expressing Csf1r (clusters 0, 1, 4, and 7) were merged and identified as MoMa; cells expressing Cd3e (clusters 2 and 8) were merged and identified as T cells; cells expressing Csf3r (cluster 3) were identified as Granulocytes (Granus); cells expressing Ube2c (clusters 5 and 9) were merged and identified as Cycling; cells expressing Ncr1 (cluster 6) were identified as Natural Killer (NK) cells; cells expressing Flt3 (cluster 10) were identified as Dendritic cells (DCs); and cells expressing CD79b (cluster 11) were identified as B cells. Right, UMAP showing the distribution of these populations at resolution 0.1 (top) and following reclassification (bottom). **C)** UMAP visualization of the tumor immune infiltrate (CD45+ cells) in individual mice. **D)** Individual cell enrichment of indicated signatures. Upper panels, UMAP visualization of the signature expression in the same UMAP plot as in Fig 1B. Lower panels, violin plots representing the signature expression by cluster. **E)** CD64^dtr^ WT or KO tumor-bearing mice were treated intraperitoneally with four doses of hDT(1ug/injection) or PBS (control) every other day, starting when tumor volume was between 30-10mm^3^. Left, schematic representation of the experimental workflow. Right, individual tumor growth curves. **F)** Dotplot illustrating TLR3 expression among myeloid subsets in four ATRT-MYC tumors. **G)** Tumor-bearing mice were treated intratumorally with three doses of PIC (50ug/injection) or PBS (control) every other day, starting when tumor volume was between 30-10mm^3^. Top, schematic representation of the experimental workflow. Bottom, individual tumor growth curves of C57BL/6J (BL/6J) tumor-bearing mice treated with PIC or PBS as control. Representative of two independent experiments. **H)** FACs-sorted live cells from R26S2 tumors were analyzed using scRNAseq (10X technology). After filtering the scRNAseq data to exclude dead, low-quality cells, immune cells (Ptprc+) and other stroma cells, we obtained 2385 tumoral cells characterized by a low/negative expression of *Smarcb1*. Top, UMAP visualization. Bottom, dot plot illustrating the scaled expression of genes associated to tumoral cells (*Ly96, Fn1, Serpine2, Acta2, Hopx* and *Smacb1)*, proliferation (*Mki67*), and *Tlr3*. **I)** C57BL/6J tumor-bearing mice were treated with three doses of intratumoral PIC (50ug/injection), three doses of intramuscular PIC (100ug/injection) or PBS (control) every other day, starting when tumor volume was between 30-10mm^3^. Tumor growth curves from RT-bearing mice treated or non-treated are presented as the mean of at least 5 mice per group, followed by individual tumor growth curves. Data shown are representative of one of two independent experiments. **J-K)** Tumor-bearing mice were treated intratumorally with three doses of PIC (50ug/injection) or PBS (control) every other day, starting when tumor volume was between 30-10mm^3^. **J)** Left, schematic representation of the experimental workflow. Right, individual tumor growth curves of TLR3 WT or KO tumor-bearing mice treated with PIC or PBS as control. Representative of two independent experiments. **K)** Left, schematic representation of the experimental workflow. Right, individual tumor growth curves of MAVS WT or KO tumor-bearing mice treated with PIC or PBS as control. Representative of two independent experiments.

**Supplementary Figure 2:**
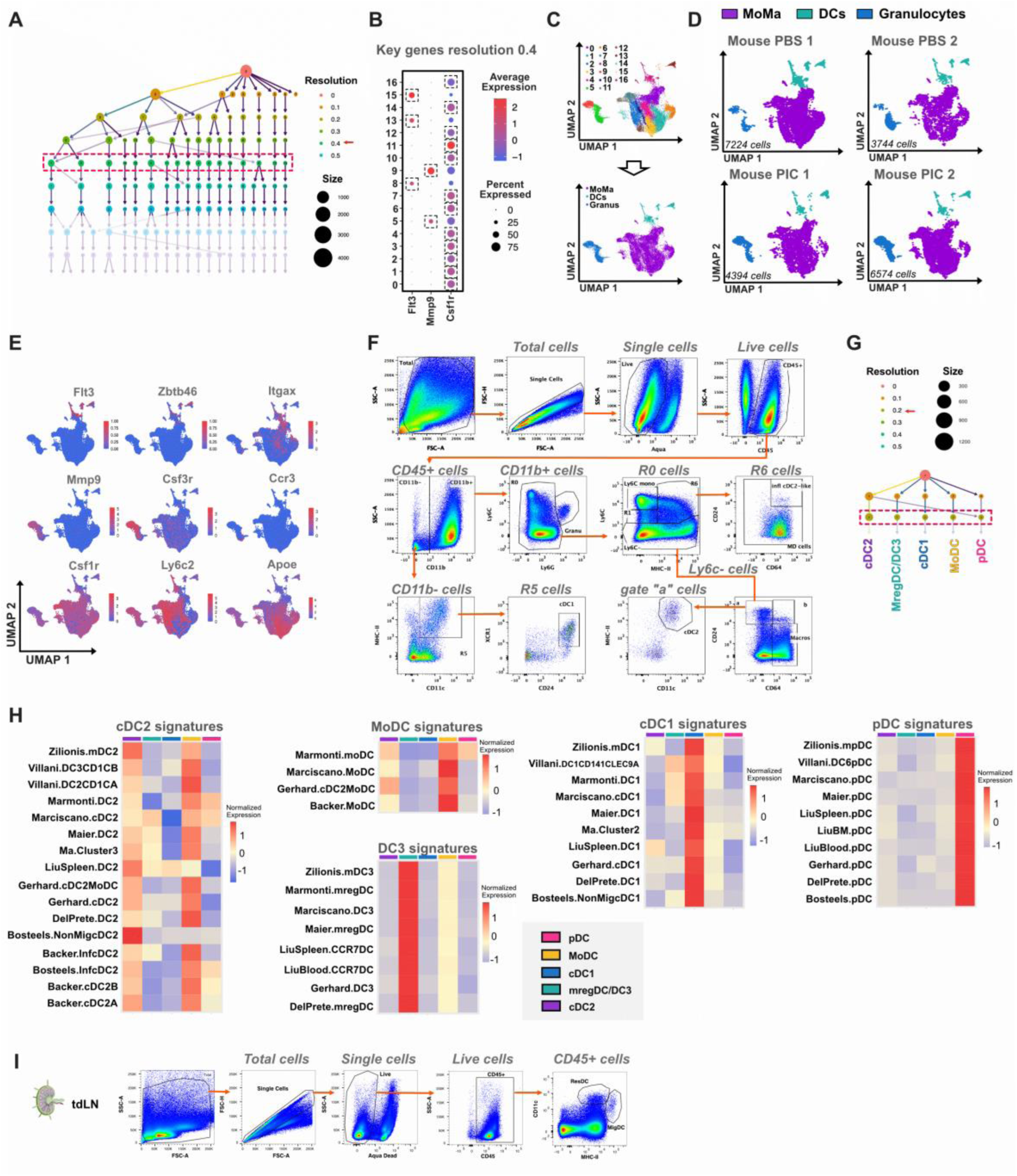
(A)Clustree visualization of scRNA-seq data showing the hierarchical relationships between cell clusters across different resolution parameters. The dotted red rectangle highlights the selected resolution. **(B)** Dot plot showing the expression of key genes used for cell clustering, with dot size representing the percentage of cells expressing each gene and color indicating scaled expression levels. The squares show the clusters that were re-grouped according to the expression of *Flt3, Mmp9* and *Cs1r*. **(C)** UMAP visualization of the resolution 0.4 containing seventeen clusters and UMAP visualization of myeloid populations after merge, containing three clusters; MoMA, granulocyte, and DC. **(D)** UMAP representation of myeloid cell distribution in each individual mouse. **(E)** Feature plots displaying the expression of key marker genes across the myeloid cell populations. **(F)** Gating strategy used for FACS-based quantification of myeloid subsets, including granulocytes, DCs, macrophages, monocyte-derived cells, and Ly6C⁺ monocytes. **(G)** Clustree visualization of scRNA-seq dendritic cell subsets data highlighting five distinct clusters: cDC2, Mreg/DC3, cDC1, MoDC, and pDC. **(H)** Heatmaps showing the normalized expression of gene signatures previously published associated with our DC clusters. **(I)** Gating strategy for FACS-based quantification of dendritic cells in tumor-draining lymph nodes (tdLN) showing resident and migratory DCs.

**Supplementary Figure 3:**
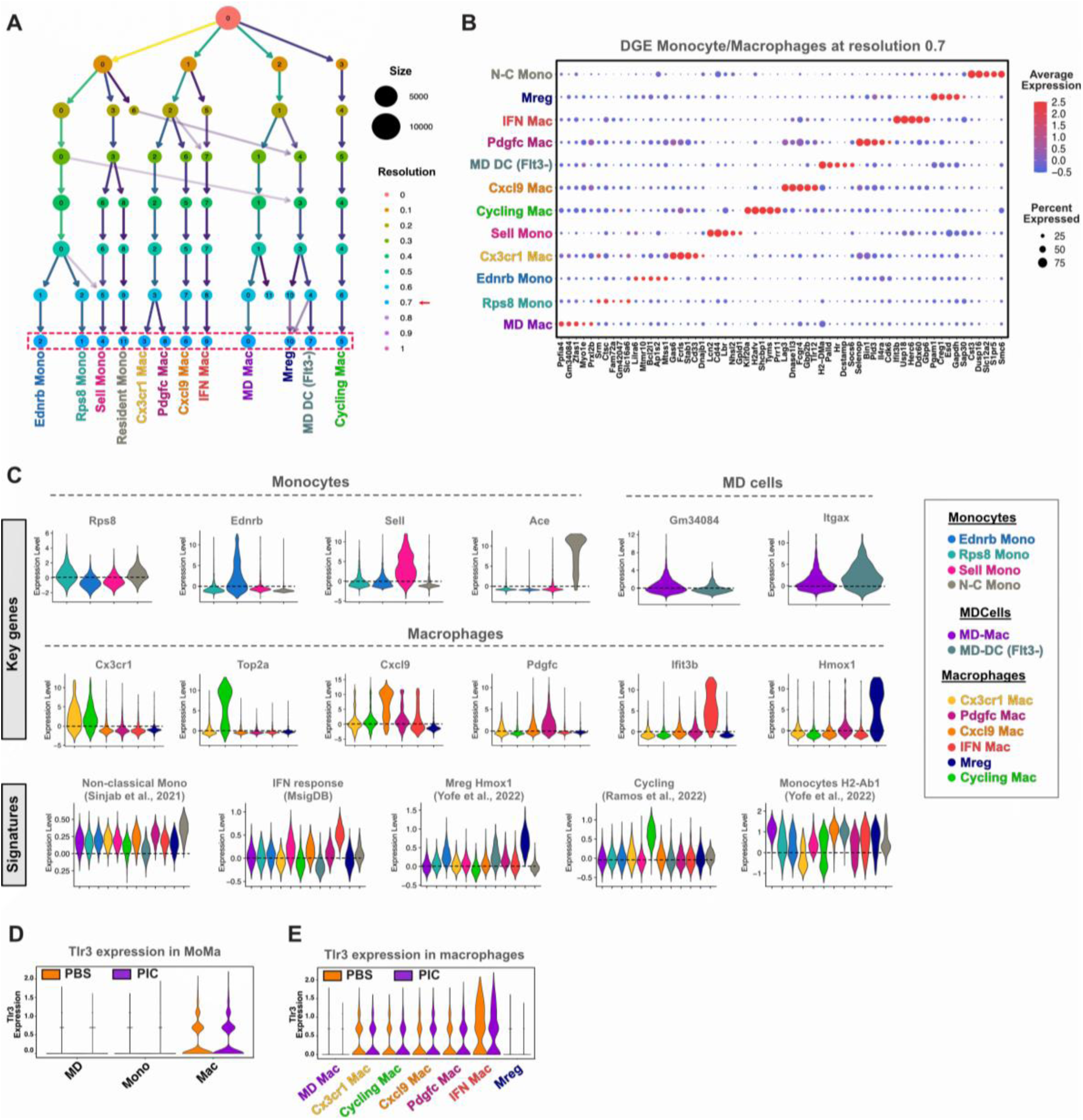
(**A**) Clustree visualization of MoMa scRNA-seq data showing the hierarchical relationships between cell clusters across different resolution parameters. The dotted red rectangle highlights the selected resolution. **(B)** Dot plot showing the scaled expression of key genes per MoMa cluster. Dot size represents the percentage of cells within each cluster expressing the corresponding gene. **(C)** Violin plots showing key gene expression for each cluster (Top) and signatures used to designate specific clusters (bottom). **(D)** Violin plot comparing Tlr3 expression levels across MoMa populations in PBS (orange) and PIC-treated tumors (violet). **(E)** Violin plot comparing Tlr3 expression levels across Macrophages subsets in PBS (orange) and PIC-treated tumors (violet).

**Supplementary figure 4:**
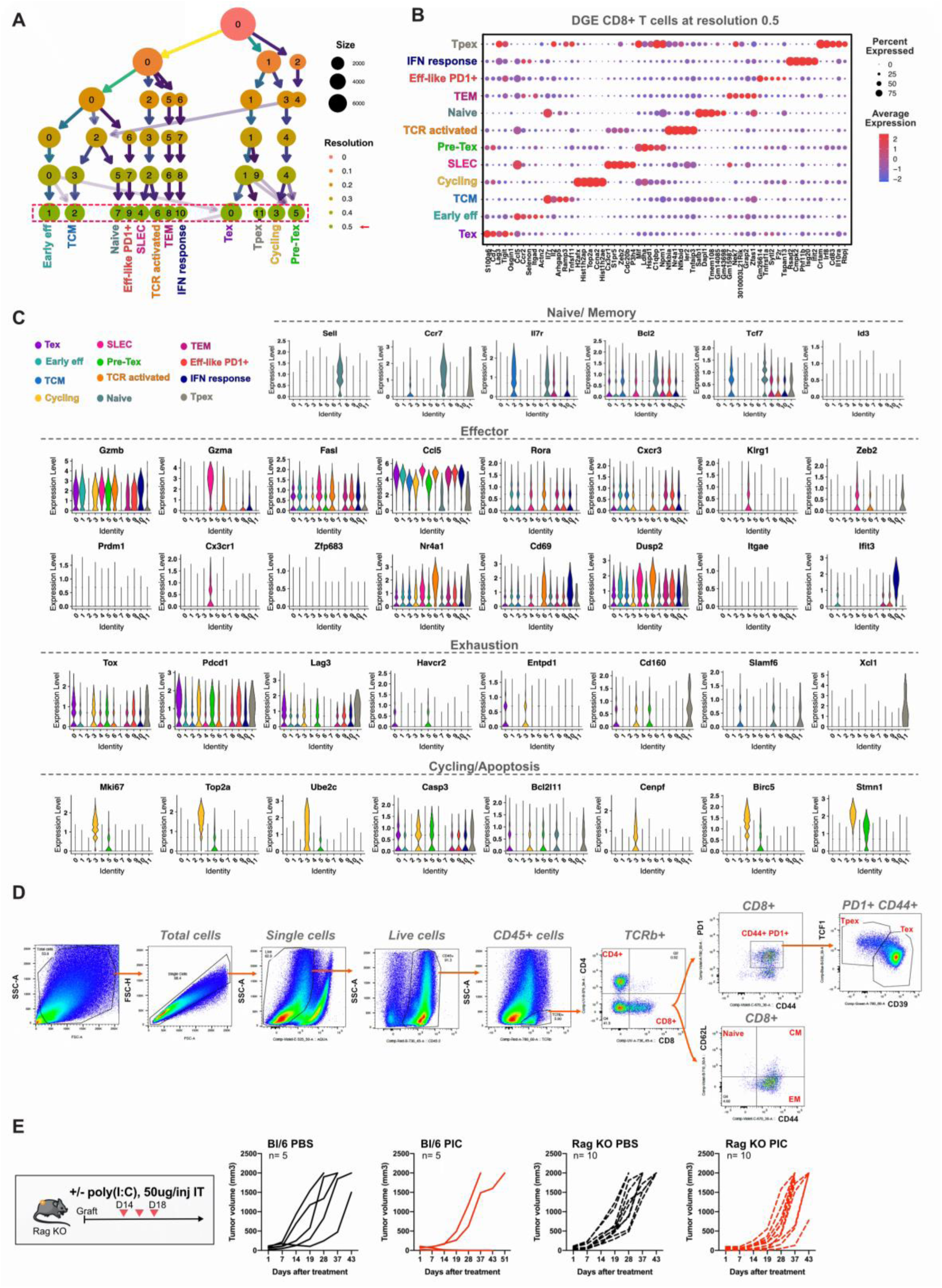
(**A**) Clustree visualization of CD8+ scRNA-seq data showing the hierarchical relationships between cell clusters across different resolution parameters. The dotted rectangle highlights the selected resolution (resolution 0.5). **(B)** Dot plot showing differential expression of key genes across CD8⁺ T cell subsets at resolution 0.5. Dot size represents the percentage of cells within each cluster expressing the corresponding gene. **(C)** Violin plots of representative genes associated with naïve/memory, effector, exhaustion, and cycling/apoptotic states. **(D)** Gating strategy for FACS-based quantification of CD8+ T cells subpopulations in tumor. **(E)** C57Bl/6 or Rag-KO tumor-bearing mice were treated intratumorally with three doses of PIC (50ug/injection) or PBS (control) every other day, starting when tumor volume was between 30-10mm^3^. Left, schematic representation of the experimental workflow. Right, individual tumor growth curves. Data shown are representative of two independent experiments, with at least n=5 mice per condition in the experiment presented and n=5 mice per condition in the replicate experiment, both yielding consisting results.

**Supplementary Figure 5:**
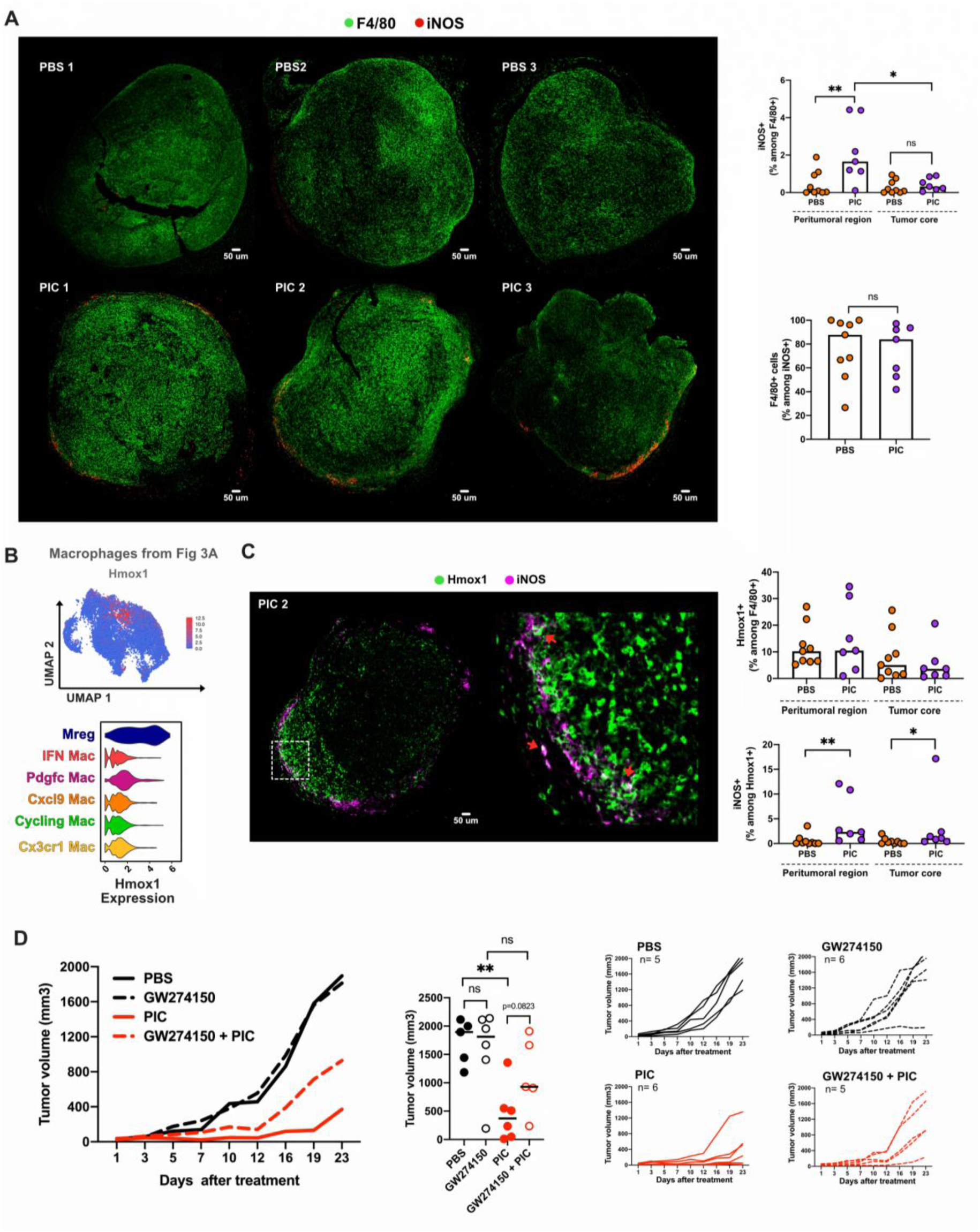
(**A**) Left: representative MICSSS images of tumor sections stained for F4/80 (green) and iNOS (red). Top panels show PBS-treated tumors (n=9), bottom panels show PIC-treated tumors (n=7). Scale bar, 50μm. Right: quantification of the percentage of F4/80+ macrophages expressing iNOS (top) and the percentage of iNOS+ cells that are F4/80+ macrophages (bottom). The peritumoral region was delineated by selecting the first 500 µm from the tumor edge. Data representative of at least 7 mice per group. **(B)** *Hmox1* expression in TAMs (UMAP from figure 3A). Top: Feature plot showing *Hmox1* expression. Bottom: Violin plots showing *Hmox1* expression among TAMs clusters. **(C)** Left, Multiplex immunohistochemical images of tumor sections from mice treated with PIC, stained with Hmox1 (green) and iNOS (magenta). Zoom-in panel (white square) at right. Scale bar: 50 μm. Right, quantification of the percentage of Hmox1+ cellos among F4/80+ macrophages (top) and the percentage of iNOS+ cells that are Hmox1+ cells (bottom). The peritumoral region was delineated by selecting the first 500 µm from the tumor edge. Data representative of at least 7 mice per group. **(D)** Tumor-bearing mice were treated intraperitoneally with seven doses of GW274150 (daily during 7 days, 100ug/injection), three intratumoral injections of PIC (every other day, 50ug/injection), with the combination of both or PBS as control, starting when tumor volume was between 30-10mm^3^ (as in Figure 5E). Left, mean tumor growth curves in each condition. Center, Tumor volume at sacrifice, day 23. Mann-Whitney test. *p<0.05, **p<0.01.

**Supplementary figure 6.**
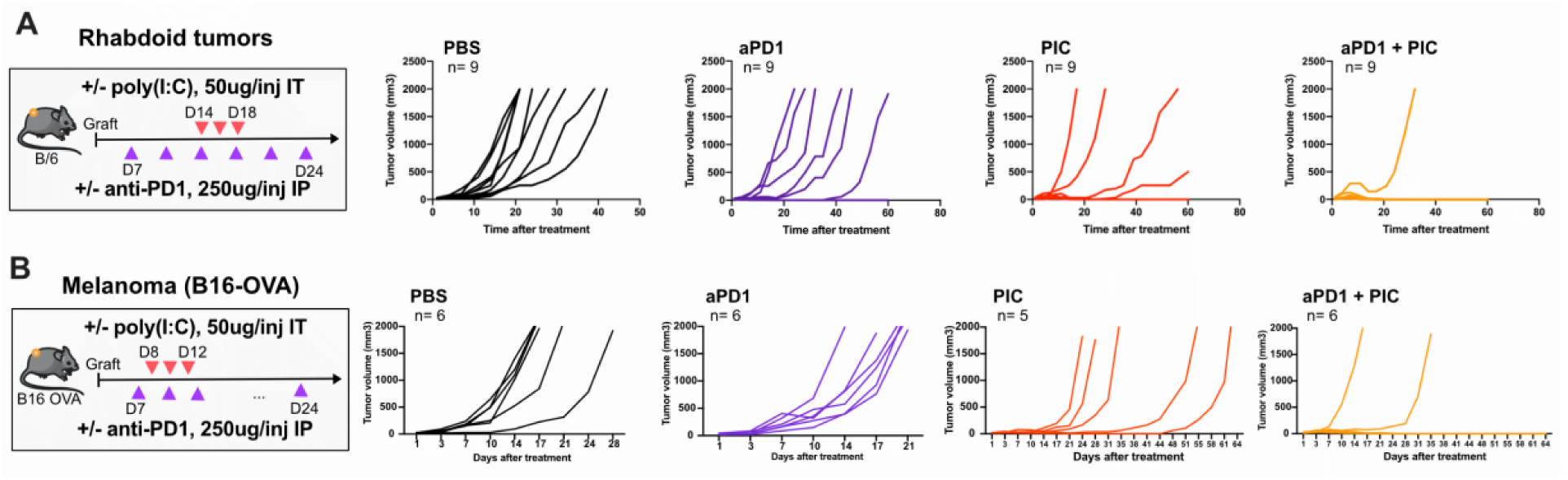
(**A**) Left, Schematic of the experimental workflow. Rhabdoid tumor-bearing mice with tumors measuring 2–4 mm in diameter (Day 7 post-engraftment) received three intratumoral injections of PIC (50 µg per dose), six intraperitoneal injections of anti–PD-1 antibody (aPD-1, 250 µg per dose) administered twice weekly, or the combination of both treatments (Combo). Control mice received PBS. Right, individual tumor growth curves. Data shown are representative of two independent experiment, with at least n=9 mice per condition. **(B)** Left, Schematic of the treatment regimen in the B16-OVA melanoma model. Mice bearing subcutaneous B16-OVA tumors (20–100 mm³) received three intratumoral injections of PIC (50 µg/injection), and/or two intraperitoneal injections per week of aPD-1 (250 µg/injection), starting one day prior to the first PIC dose. Control mice received PBS. Right, individual tumor growth curves.

**Supplementary figure 7:**
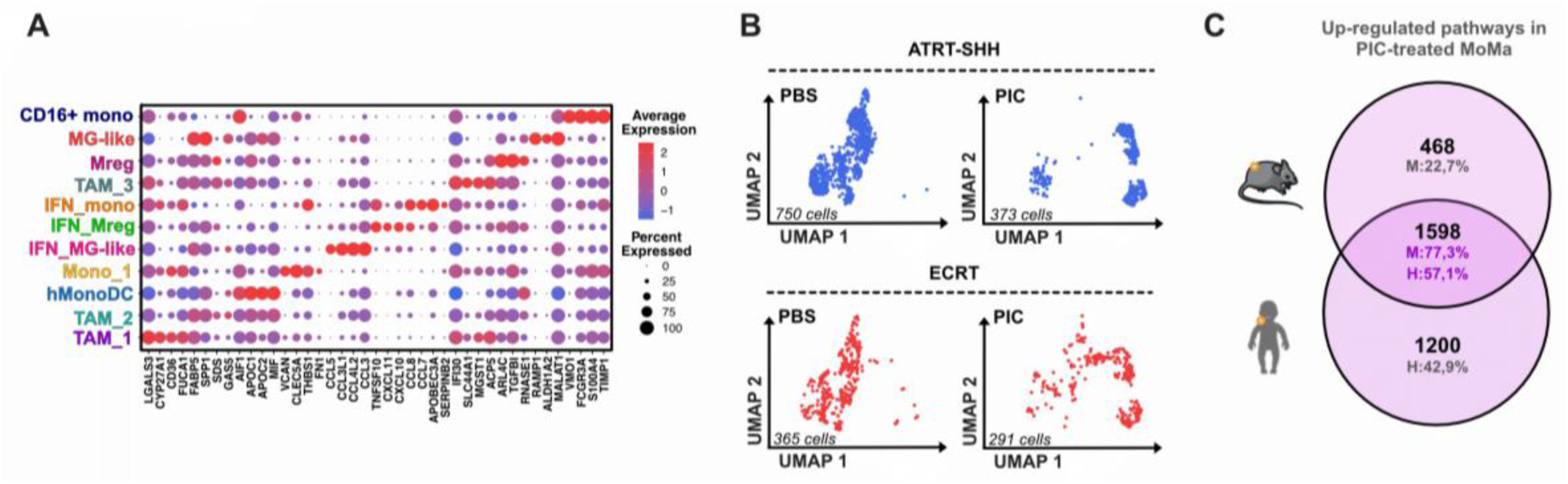
(**A**) Dot plot showing the expression of representative lineage and functional gene signatures across identified MoMa subclusters. Dot size indicates the percentage of cells within each cluster expressing a given gene, and color scale represents average normalized expression. **(B)** UMAP plots of MoMa cells isolated from ATRT-SHH (top) and ECRT (bottom) tumors after 16h ex vivo culture with PBS or PIC (50 µg/mL). Cell numbers per condition are indicated. **(C)** Venn diagram showing the number of overlapping up-regulated pathways following PIC treatment in mouse (M) and human (H) MoMa.

